# Ferroptotic stress promotes the accumulation of pro-inflammatory proximal tubular cells in maladaptive renal repair

**DOI:** 10.1101/2021.03.23.436661

**Authors:** Shintaro Ide, Yoshihiko Kobayashi, Kana Ide, Sarah A. Strausser, Savannah Herbek, Lori L. O’Brien, Steven D. Crowley, Laura Barisoni, Aleksandra Tata, Purushothama Rao Tata, Tomokazu Souma

## Abstract

Overwhelming lipid peroxidation induces ferroptotic stress and ferroptosis, a non-apoptotic form of regulated cell death that has been implicated in maladaptive renal repair in mice and humans. Using single-cell transcriptomic and mouse genetic approaches, we show that proximal tubular (PT) cells develop a molecularly distinct, pro-inflammatory state following injury. While these inflammatory PT cells transiently appear after mild injury and return to their original state without inducing fibrosis, they accumulate and contribute to persistent inflammation after severe injury. This transient inflammatory PT state significantly downregulates glutathione metabolism genes, making them vulnerable to ferroptotic stress. Genetic induction of high ferroptotic stress in these cells after mild injury leads to the accumulation of the inflammatory PT cells, enhancing inflammation and fibrosis. Our study broadens the roles of ferroptotic stress from being a trigger of regulated cell death to include the promotion and accumulation of proinflammatory cells that underlie maladaptive repair.

## Introduction

Acute kidney injury (AKI) afflicts 1.2 million hospitalized patients annually in the US; twenty to fifty percent of AKI survivors progress to chronic kidney disease (CKD), increasing their risk for dialysis-dependency, cardiovascular events, and mortality (1–3). Other than general supportive care, there are no targeted therapies to treat AKI or to prevent AKI to CKD transition. Understanding the molecular events underpinning the AKI to CKD transition is needed to develop therapeutic strategies to interrupt this devastating disease process.

Clinical and preclinical studies have identified damage to proximal tubular (PT) epithelial cells after severe AKI as a critical mechanism driving transition to CKD (3–8). PT cells are most severely affected by acute ischemic and toxic injuries due to their high metabolic and energy-intensive transporter activities required to maintain normal homeostasis of body fluids (7–9). In the renal repair process, damaged PT cells adopt heterogeneous molecular states (10). They reactivate genes normally active during renal development (11–13), alter their dependency on metabolic fuels (14), change their morphology, and proliferate to replenish the areas of denuded epithelium in the proximal tubule (7, 15). When the initial damage to kidneys is mild, PT cells subsequently return to their original state by redifferentiation with resolution of inflammation and fibrosis (7, 10, 11, 14–17). However, if damage is more extensive, prolonged, or recurrent, the damaged cells fail to redifferentiate, leading to persistent inflammation, fibrosis, and eventual cell death. The molecular pathways that govern proximal tubular heterogeneity and cell fate during failed renal repair after severe injury are poorly understood. This knowledge gap prevents the development of therapies based on underlying disease mechanisms.

One of the critical pathways involved in AKI pathogenesis and proximal tubular cell death is ferroptosis, a distinct non-apoptotic form of regulated cell death (18–25). An imbalance between the generation of lipid peroxides and their detoxification induces overwhelming accumulation of lipid peroxides (ferroptotic stress), triggering ferroptosis (18, 23). Glutathione/glutathione peroxidase 4 (GPX4) axis is the central defense pathway to prevent ferroptotic stress and ferroptosis (18–20, 26). Global genetic deletion of *Gpx4* causes renal tubular epithelial death and acute kidney injury, identifying renal tubular epithelial cells as one of the most vulnerable cell types to ferroptotic stress (26). Moreover, reduced glutathione and NADPH availability further render ischemia-reperfusion injured kidneys vulnerable to ferroptotic stress (3, 27). Accumulating evidence suggests that pharmacological inhibition of ferroptotic cell death ameliorates AKI severity and excess ferroptotic stress has been linked to failed renal repair in patients, suggesting a new therapeutic target (24–26, 28). However, it is still not clear whether ferroptotic stress has additional roles in the pathogenesis of AKI and its sequelae beyond the induction of cell death and loss of functional tubular cells.

Here, using complementary single-cell transcriptomic and mouse genetic approaches, we identify the role of a molecularly distinct damage-associated PT cell state that is dynamically and differentially regulated during successful versus failed repair. Furthermore, we provide mechanistic evidence that ferroptotic stress in PT cells enhances this damage-associated state, thereby promoting failed renal repair and the AKI-to-CKD transition.

## Results

### Tubular epithelial cells exhibit heterogeneous molecular states after severe injury

To identify cellular mechanisms that promote maladaptive repair after severe kidney injury, we first developed and optimized mouse models for “successful” versus “failed” renal repair after ischemia-reperfusion-induced injury (IRI). This was achieved by extending renal ischemic times from 20 min for successful recovery to 30 min for failed recovery (Fig. S1 and S2). After mild injury, histologic examination showed that inflammation and macrophage accumulation resolved within 21 days (ischemic time 20 min; Fig. S1, D and E). By contrast, after severe injury there was progressive epithelial damage and fibrosis, and the accumulation of F4/80+ macrophages persisted around the damaged epithelial cells for at least 6 months (ischemic time 30 min; Fig. S1, D and E; and S2E).

We used this failure-to-repair model (unilateral IRI, ischemic time 30 min) to generate a single-cell transcriptome map of failed renal repair (Fig. 1A). Kidneys were harvested at 6 hours and 1, 7, and 21 days after IRI. High-quality transcriptome data from a total of 18,258 cells from injured kidneys (IRI) and homeostatic uninjured kidneys (Homeo) were obtained (Fig. 1, B and C). Using a Seurat integration algorithm that normalizes data and removes potential batch effects (29, 30), we integrated the transcriptome data from each condition and performed unsupervised clustering analysis of the integrated dataset. Uniform manifold approximation and projection (UMAP) resolved 21 separate clusters, representing distinct cell types (Fig. 1B; Fig. S3B and S4A). The cellular identity of each cluster was determined based on known cell type-specific markers (31, 32). We successfully identified known cell-type-specific damage-induced genes such as *Havcr1* (kidney injury molecule-1, KIM1), *Krt8* (keratin 8), *Krt20* (keratin 20), and *Lcn2* (neutrophil gelatinase-associated lipocalin, NGAL) selectively in ischemia-reperfusion-injured (IRI) kidneys, but not in homeostatic uninjured control kidneys (Fig. S3C), (5, 33, 34).

**Figure 1.**
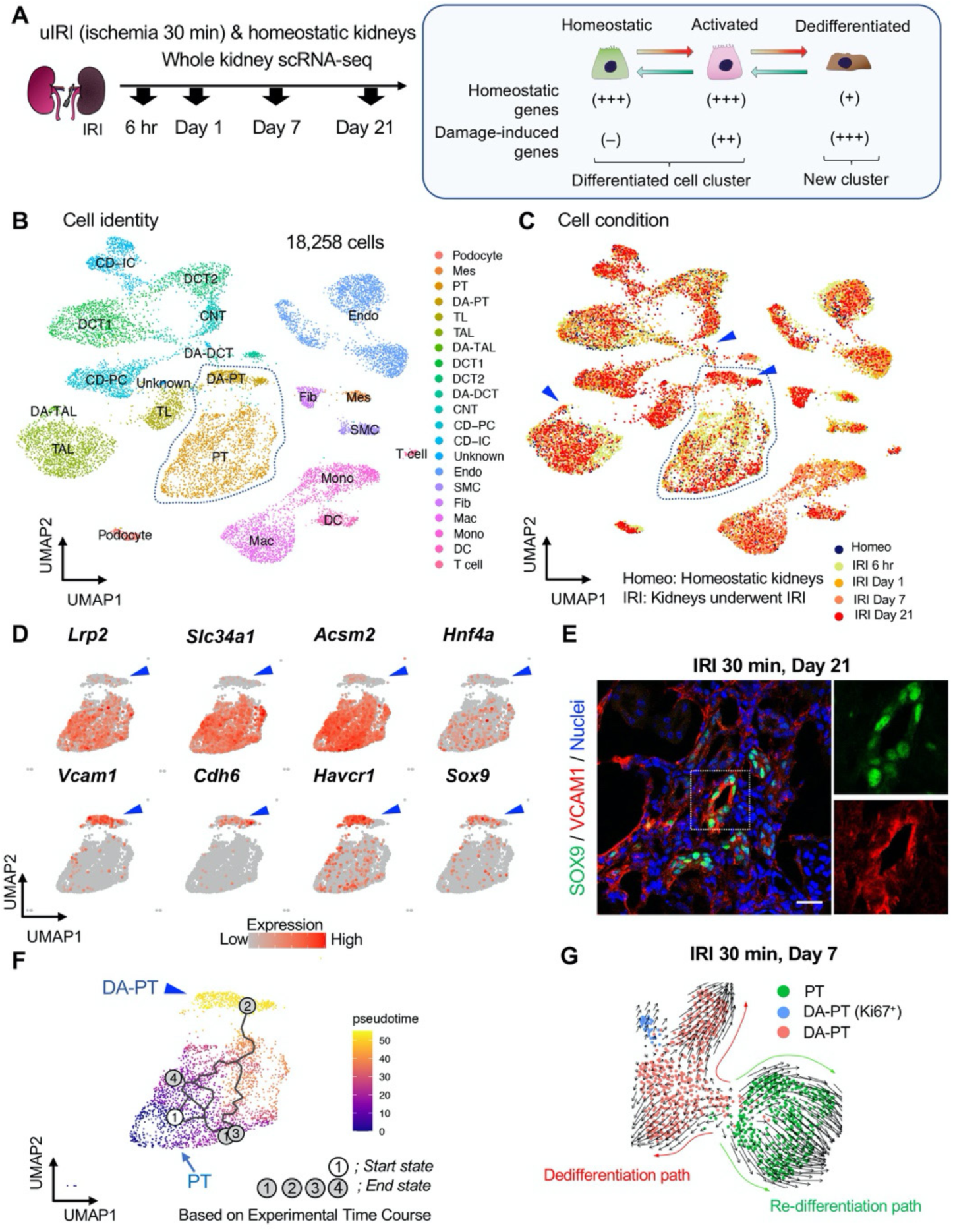
single-cell RNA sequencing (scRNA-seq) identifies dynamic cellular state transitions of tubular epithelial cells after severe IRI. (A) Drop-seq strategy. uIRI, unilateral IRI. A schematic illustration of epithelial cell states is shown. (B) and (C) Integrated single-cell transcriptome map. Unsupervised clustering identified 21 distinct clusters in the UMAP plot. Arrowheads indicate damage-associated tubular epithelial cells. The dotted area (PT cell clusters; PT and DA-PT) was used for the downstream analyses in (D)–(G). (D) UMAP plots showing the expression of indicated genes in PT cell clusters (PT and DA-PT in (B)). Differentiated/mature PT cell markers: *Lrp2* (megalin), *Slc34a1* (sodium-dependent phosphate transporter 2a, NaPi2a), *Acsm2* (Acyl-coenzyme A synthetase), and *Hnf4a* (hepatocyte nuclear factor 4*α*); and damage-induced genes: *Vcam1* (vascular adhesion molecule 1), *Cdh6* (Cadherin 6), *Havcr1* (kidney injury molecule-1, KIM1), *Sox9* (Sry-box 9). Arrowheads; DA-PT. (E) Immunostaining for SOX9 and VCAM1 using post-severe IRI kidneys on day 21. Scale bar: 20 μm. (F) Pseudotime trajectory analysis of proximal tubular cells (PT and DA-PT clusters) that underwent IRI. A region occupied with cells from 6-hrs after post-IRI was set as a starting state. (G) RNA velocity analysis of PT clusters (PT and DA-PT) from post-IRI kidneys on day 7. Cells in PT clusters from IRI day 7 dataset was extracted for the analysis. The arrows indicate predicted lineage trajectories. Abbrev: PT, proximal tubule; DA-PT, damage-associated PT; TL, thin limb; TAL, thick ascending limb; DA-TAL, damage-associated TAL; DCT, distal convoluted tubule; DA-DCT, damage-associated DCT; CNT, connecting tubule; CD, collecting duct (P, principal cells, IC, intercalated cells); Mes, mesangial cells; Endo, endothelial cells; SMC, smooth muscle cells; Fib, fibroblasts; Mac, macrophages; Mono, monocytes; DC, dendritic cells.

Based on the cell clustering and gene expression patterns, we noticed that there are at least three epithelial cell states (homeostatic normal, activated, and dedifferentiated cells) in our dataset (See Fig. 1A, right panel). Homeostatic normal cells express high expression of “anchor” genes involved in normal cell function and identity (Fig. S3, B and C). Most of the tubular epithelial cells from IRI kidneys robustly expressed damage-induced genes (ex. *Havcr1*, *Krt8*, *Krt20*, *Lcn2*), indicating they are in activated states (Fig. S3C). These activated cells and homeostatic cells were grouped in the same cluster because they both highly express anchor genes characteristic for normal tubular epithelial states and functions (Fig. 1, B and C; Fig, S3C). However, we also identified additional damage-associated tubular epithelial clusters (Fig. 1C, arrowheads; DA-PT, DA-TAL, and DA-DCT) that had lost or reduced expression of “normal” mature epithelial cell marker genes but highly expressed damage-induced genes (Fig. S3, B and C, Fig. 1A).

Among these damage-associated epithelial cell clusters, we found a damage-associated proximal tubular cell state (See DA-PT cluster), which shows reduced homeostatic gene expression (ex. *Lrp2*, *Slc34a1*, *Hnf4a*, and *Acsm2*) and enrichment for genes associated with both renal development and kidney injury in human and mouse (ex. *Cdh6*, *Sox9*, *Sox4*, *Cited2*, *Vcam1*, *Vim*, and *Havcr1*; Fig. 1D, and Fig. S5B and S6A), (35–37). Moreover, gene ontology enrichment analyses of this cellular population revealed proinflammatory molecular signatures and enriched expression of chemokines and cytokines such as *Cxcl2, Cxcl1, Ccl2*, and *Spp1* (Fig. S5B, D, and E). Reduced expression of *Hnf4a*, which is a transcription factor essential for the maturation of PT cells (38, 39), and other homeostatic genes and upregulation of *Cdh6*, which is selectively expressed in immature proximal tubule progenitors in development and is essential for renal epithelialization (39–41), suggest that the cells in this cluster (DA-PT) are in a less differentiated cell state (Fig. 1D and Fig. S6A), (39). Then, we compared the transcriptional signature of this damage-associated PT cell state with previously published neonatal kidney single-cell RNA seq data (GSE94333, Fig. S7, A and B), (37). The top 100 genes enriched in immature early PT cells in neonatal kidneys were mainly expressed in this damage-associated PT cell state (DA-PT, Fig. S7, C and D). These analyses support our notion that the cells in the DA-PT cluster are in a dedifferentiated inflammatory state.

Among the damage-induced genes expressed in this dedifferentiated inflammatory PT cell state, we focused on the enrichment of *Sox9* and *Vcam1* (Fig. 1D, See DA-PT cluster, arrowheads). Recent single-nucleus transcriptomic profiling of mouse IRI-kidneys identified vascular cell adhesion molecule 1 (VCAM1) as a marker of non-repairing proximal tubular cell state (10), and *Vcam1* induction has been observed in multiple forms of human kidney diseases, including allograft rejection (42). SRY-box9 (SOX9) is an essential transcription factor for successful renal repair after acute ischemic and toxic insults (12, 13) and is involved in the development of multiple organs, including mouse and human kidneys (43). SOX9 contributes to tissue repair processes by conferring stemness, plasticity, and regenerative capacity (12, 13, 44–47). Our single-cell RNA-sequencing (scRNA-seq) data revealed that *Sox9* was most robustly induced in damage-associated PT cells compared to other tubular epithelial cells (DA-PT, Fig. S5C). To validate this finding, we performed immunofluorescence for SOX9 and VCAM1 in histological sections of kidneys with failed repair. SOX9 nuclear accumulation was observed in VCAM1^+^ proximal tubular cells (Fig. 1E). High expression of *Sox9* and *Vcam1* suggests a potential role of this damage-associated PT cell state both in adaptive and maladaptive renal repair in a context-dependent manner, such as ranging severity of injury.

To understand the lineage hierarchy of PT cell states, we analyzed the PT cells from differentiated and damage-associated PT cell clusters (PT and DA-PT in Fig. 1B) using two computational algorithm tools (Monocle 3 and Velocyto) that allow the prediction of cell differentiation trajectories (48, 49). We observed a predicted differentiation trajectory originating from PT to DA-PT by placing each cell from the entire dataset in pseudotime (Fig. 1F and Fig. S8, A and B). We then performed RNA velocity analysis, which predict the cell state trajectory based on the ratio between unspliced and spliced mRNA expressions, for these two PT cell states from the post-IRI dataset on day 7. Our RNA velocity analysis showed two trajectories running in opposite directions from the middle of the cluster, a position where genes associated with tubular maturation and damage are both not highly expressed (Fig. 1G and Fig. S8C). One projects towards the area with high damage-induced genes (dedifferentiation path to damage-associated PT cell state) and the other towards the area with high maturation-associated genes (redifferentiation path to differentiated PT cell state), suggesting the existence of cellular plasticity at this stage (Day 7 post-IRI; Fig. 1G and Fig. S8C).

### Proximal tubular cells dynamically alter their cellular states after acute kidney injury

To determine the temporal dynamics of damage-associated PT cell state in adaptive and maladaptive repair, we performed expression analyses of marker genes for this PT cell state in successful and failed renal repair processes. Quantitative RT-PCR analyses for *Sox9*, *Cdh6*, and *Vcam1* genes confirmed the transient induction of these genes and resolution after mild ischemic injury (20 min ischemia, Fig, 2B), but persistently elevated expression after severe ischemic injury (30 min) through 21 days after injury (Fig. 2B). Using immunofluorescence and *in situ* hybridization, we observed more VCAM1^+^ and *Cdh6*^+^ tubular epithelial cells in IRI kidneys after 30 min than 20 min ischemia (Fig. 2C). The number of SOX9 positive cells was similarly increased between kidneys with mild and severe IRI on day 7 compared to baseline (Fig. 2, D and E). Confocal imaging showed that most of the SOX9-positive cells co-express VCAM1 (Fig. S9, B and C). Notably, SOX9 expression was reduced to baseline level in the kidneys that underwent mild injury while it persisted up to 6 months after severe IRI (Fig. 2, D and E). We observed clusters of SOX9^+^VCAM1^+^ cells in the remaining parenchyma at 6 months post-severe IRI (Fig. 2F and Fig. S9, B and C). In accordance with the hypothesis that severe IRI injury is associated with increased signature of damage-associated PT cell state, there was a reduction of homeostatic gene expressions (*Acsm2* and *Slc34a1*) and the number of fully differentiated PT cells, which have high lotus tetragonolobus lectin (LTL)-binding (Fig. 2B and Fig. S6, B–D), (39). This finding is in line with a clinical correlation between low expression of *ACSM2B*, the human ortholog of *Acsm2*, and reduced renal function in patients with CKD (50). Collectively, these data support the emergence and accumulation of damage-associated PT cells after severe injury but their return to a homeostatic state after mild injury.

**Figure 2.**
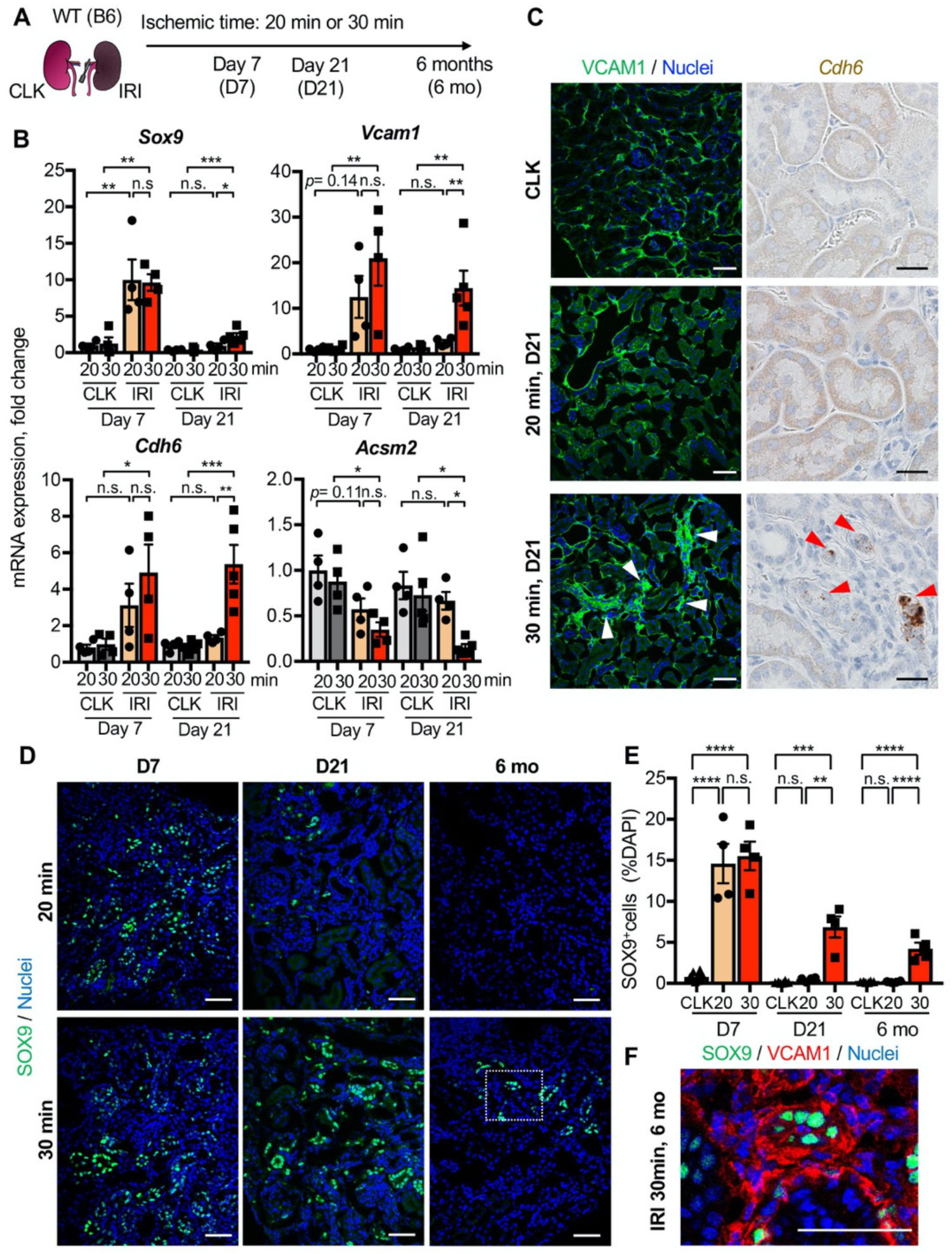
Damage-associated PT cells emerge transiently after mild injury but persist after severe injury. (**A**) Experimental workflow for the mild and severe IRI models. Left kidneys from wild-type (WT) C57BL/6J (B6) mice were subjected to mild (20 min) and severe (30 min) ischemia (unilateral IRI, uIRI). Contralateral kidneys (CLK) were used as controls. (**B**) Real-time PCR analyses of indicated gene expression. Whole kidney lysates were used. N = 4-5. (**C**) Expression analyses of VCAM1 and *Cdh6*. Immunostaining for VCAM1 revealed clusters of VCAM1^high^ tubular epithelial cells. *In situ* hybridization (ISH) was used to detect *Cdh6* gene expression on kidney sections. (**D**) Immunostaining for SOX9 in mild (20 min) and severe (30 min) IRI kidneys collected at indicated time points. **(E)** Quantification of SOX9^+^ cells over the DAPI^+^ area. Note that SOX9^+^ cells persist after severe IRI up to 6 months after IRI (30 for 30 min ischemia). In contrast, they disappear after a transient appearance in post-mild IRI kidneys (20 for 20 min ischemia). N= 4-8. (**F**) Immunostaining for SOX9 and VCAM1 (6 months post-severe IRI kidneys, dotted area in D). Scale bars, 20 μm in (C, *Cdh6*), and 50 μm (C, VCAM1, D and F). \**P* < 0.05; \*\**P* < 0.01; \*\*\**P* < 0.001; \*\*\*\**P* < 0.0001, one-way ANOVA with post hoc multiple comparisons test. Abbrev. n.s., not significant.

To further characterize the dynamic changes and plasticity of proximal tubular cell state, we employed a *CreERT2* allele of *Sox9*, a highly enriched gene in the damage-associated PT cell state (DA-PT, Fig. S5C), combined with *Rosa26^tdTomat^*° reporter to carry out lineage tracing (Fig. 3A). In this mouse line (*Sox9^IRES-CreERT2^; Rosa26^tdTomato^*), tamoxifen administration permanently labels the *Sox9*-lineage cells with the tdTomato fluorescent reporter and provides the spatial information of the cells with a history of *Sox9* expression. On day 21, we found that severe ischemia (30 min) induces more robust accumulation of *Sox9*-lineage labeled cells than mild ischemic injury (20 min) in the cortex and outer medulla of the IRI-kidneys (Fig. 3, B and C). Approximately 25 percent of *Sox9*-lineage cells that underwent severe IRI were positive for VCAM1 on day 21, suggesting that part of *Sox9*-lineage cells did not fully redifferentiate after severe injury (Fig. 3, D and E; 30 min). In contrast, only a few *Sox9*-lineage cells that underwent mild injury were VCAM1 positive at this time, indicating successful redifferentiation (Fig. 3, D and E; 20 min). Taken together, our data suggest that the loss of plasticity and the impaired redifferentiation of damage-associated PT cells underlie the failed renal repair/regeneration process (Fig. 3F).

**Figure 3.**
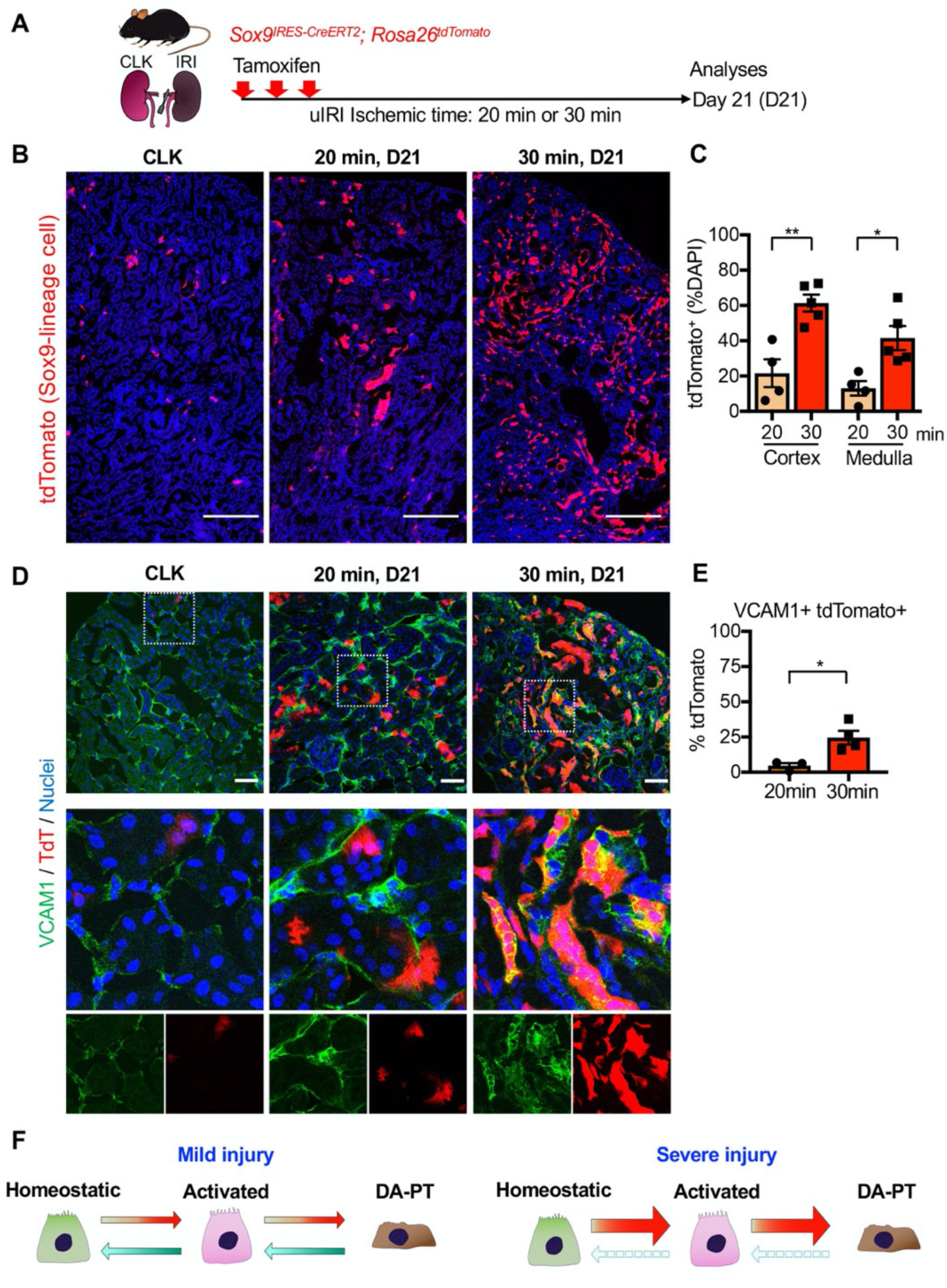
Lineage-tracing identifies the cellular plasticity of damage-associated PT cells. (**A**) Schematic of fate-mapping strategy using *Sox9^IRES-CreERT2^*; *Rosa26^tdTomato^* mice. Tamoxifen was administered 3 times on alternate days. Contralateral kidneys (CLK) were used as controls. (**B**) Distribution of tdTomato-expressing cells (*Sox9*-lineage cells) in contralateral (CLK), mild (20 min) and severe (30 min) IRI kidneys on day 21 (D21). (**C**) Quantification of tdTomato^+^ area relative to DAPI^+^ area in (B). DAPI was used for nuclear staining. N = 4-5. (**D**) Immunostaining for VCAM1 in *Sox9*-lineage-tagged kidneys. *Sox9*-lineage cells express native tdTomato red fluorescence (TdT). Insets: individual fluorescence channels. (**E**) Quantification of double-positive cells in total tdTomato^+^ cells in (D). N = 3-4. Note that more *Sox9*-lineage cells express VCAM1 after severe IRI (30 min) compared to mild IRI (20 min) on day 21. One-way ANOVA with post hoc multiple comparisons test and unpaired Student’s t-test were used for (C) and (E), respectively. Scale bars, 200 μm in (B); and 50 μm in (D). \**P* < 0.05; \*\**P* < 0.01; \*\*\**P* < 0.001; \*\*\*\**P* < 0.0001. (**F**) Schematic illustration of PT cell state dynamics. Differentiated/mature PT cells are activated, transit into a molecularly distinct PT cell state (damage-associated PT cells in DA-PT cluster), and redifferentiate into their original state after mild injury (left). Severe injury prevents the redifferentiation of damage-associated PT cells into normal PT cell state, leading to the accumulation and persistence of damage-associated PT cells (right).

### Damage-associated PT cells create a proinflammatory milieu with renal myeloid cells

While an initial inflammatory response is critical for tissue repair, uncontrolled persistent inflammation underlies organ fibrosis (7–9). We hypothesized that the accumulation of damage-associated PT cells, which show proinflammatory transcriptional signature (Fig. S5B, D, and E), creates an uncontrolled inflammatory milieu by interacting with resident and infiltrating myeloid cells such as macrophages and monocytes (51). To determine the intercellular interactions between damage-associated PT cells and myeloid cells, we used NicheNet, a computational algorithm tool that infers ligand-receptor interactions and downstream target genes (Fig. 4, A-D), (52). We applied NicheNet to predict ligand-receptor pairs in which ligands from damage-associated PT cells interact with receptors in monocyte or macrophages (Fig. 4, A and C), (52). Among the top 5 predicted ligands expressed in damage-associated PT cells, we confirmed the enrichment of *Icam1*, *Pdgfb*, and *Apoe* expression in this cell state (Fig. 4E, arrowheads). *Icam1* and *Pdgfb* have been implicated in human AKI (36), and *Apoe* genetic variation has been linked with CKD progression (53). As inferred by NicheNet, mRNA expression of *Icam1*, *Pdgfb*, and *Apoe* were markedly increased in the kidneys showing the accumulation of damage-associated PT cells compared to post-IRI kidneys without the accumulation (Fig. 4F; 30 min *vs.* 20 min ischemia). These data delineate a complex inflammatory circuit within the damaged kidneys involving intercellular communication between damage-associated PT cells and myeloid cells that contribute to maladaptive renal repair.

**Figure 4.**
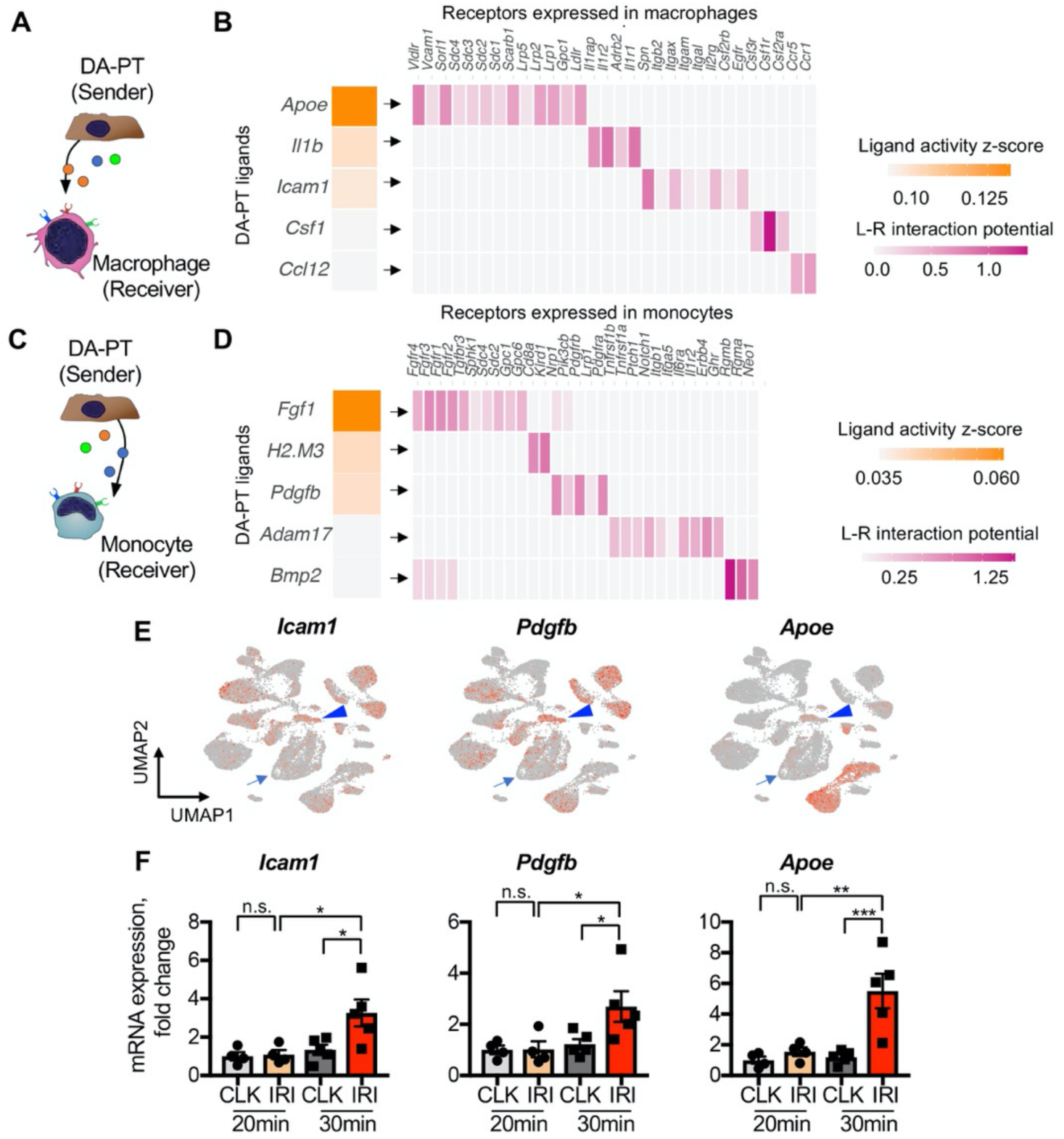
Damage-associated PT cells create a proinflammatory milieu with myeloid cells. (**A**) Schematic model of intercellular communications between damage-associated PT cells and macrophages. NicheNet was used to predict intercellular interactions using our integrated single-cell map of failed renal repair. (**B**) Predicted ligands from damage-associated PT cells and receptors in macrophages. (**C**) Schematic model of intercellular communications between damage-associated PT cells and monocytes. (**D**) Predicted ligands from damage-associated PT cells and receptors in monocytes. (**E**) UMAP plots showing the expression of indicated genes. Our integrated single-cell map of mouse failed renal repair is shown (See Fig. 1, B and C). Arrowheads indicate damage-associated PT cells (DA-PT cluster). Arrows indicate differentiated PT cells (PT cluster). (**F**) Real-time PCR analyses of indicated gene expression. Post-IRI kidneys on day 21 that underwent mild (20 min) or severe (30 min) ischemia were used. N = 4-5. \**P* < 0.05; \*\**P* < 0.01; \*\*\**P* < 0.001, one-way ANOVA with post hoc multiple comparisons test.

### Damage-associated PT cells exhibit high ferroptotic stress after severe IRI

Next, we investigated the molecular mechanisms that are critical for cells to traverse between differentiated PT cells and damage-associated PT cells. To this end, we analyzed the transcriptional signature of PT cells in the differentiated/mature cluster to identify critical pathways to maintain this cellular state. We found that genes associated with glutathione metabolic processes and anti-oxidative stress response pathways are overrepresented in the differentiated mouse PT cell cluster (Fig. 5A; Fig. S5F; S10A). We also found that these pathways are enriched in normal differentiated human PT cells (Fig. 5A and Fig. S11, A and B; GSE131882). Mirroring these findings, oxidative stress-induced signaling pathways related to failed renal repair, such as cellular senescence and DNA damage responses (54, 55), were highly enriched in damage-associated PT cells (Fig. S10A). Taken together, we propose that glutathione-mediated anti-oxidative stress responses are critical for maintaining the cellular identity of fully differentiated PT cells, and dysregulation of these pathways underlies the failure of damage-associated PT cells to redifferentiate into normal PT cell state.

**Figure 5.**
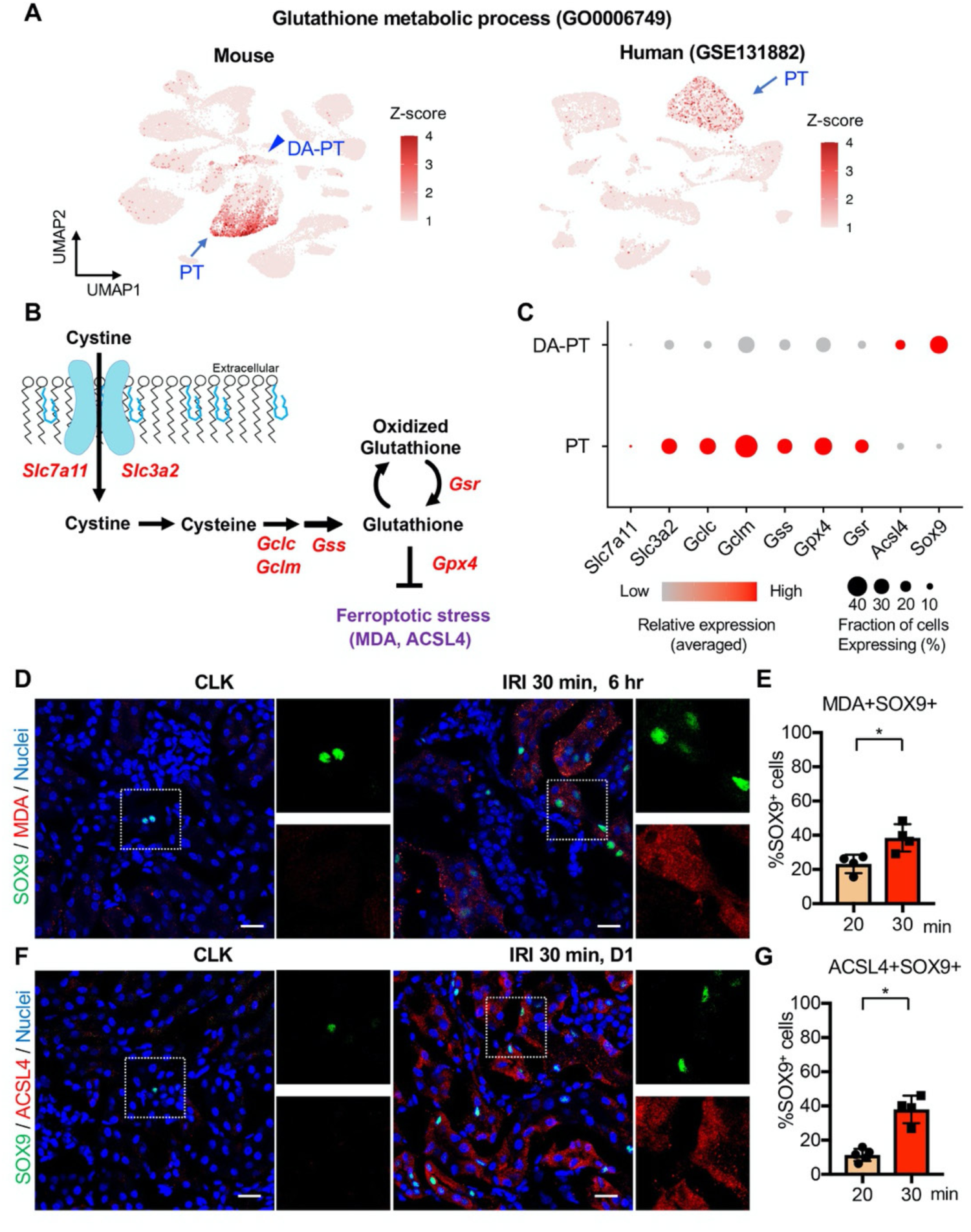
Damage-associated PT cells undergo high ferroptotic stress after severe IRI. (**A**) UMAP rendering of glutathione metabolic process in mouse and human kidneys. (**B**) A scheme showing glutathione-glutathione peroxidase 4 (GPX4) anti-ferroptotic defense pathway. *Slc7a11* and *Slc3a2* (system x_c_^-^); *Gclc* and *Gclm* (glutamate-cysteine ligase); *Gss* (glutathione synthetase); *Gsr* (glutathione reductase): and *Gpx4*. MDA (malondialdehyde, a lipid peroxidation product) and ACSL4 (acyl-CoA synthetase long-chain family member 4) are markers for ferroptotic stress. (**C**) Dot plots show the expression of genes for glutathione-GPX4 axis, *Sox9,* and *Acsl4*. (**D)** Immunostaining for SOX9 and MDA, and (**E**) quantification of double-positive cells in total SOX9^+^ cells. N = 4. (**F**) Immunostaining for SOX9 and ACSL4, and (**G**) quantification of double-positive cells in total SOX9^+^ cells. N = 4. Insets: individual fluorescence channels of the dotted box area. Note that severe ischemia (30 min) induces more ferroptotic stress markers (MDA and ACSL4) in SOX9^+^ cells in damaged kidneys than mild ischemia (20 min). Wild-type C57BL/6J mice were used for (D) to (G). Scale bars, 20 μm in (D) and (F). \**P* < 0.05. unpaired Student’s t-test.

Among the cellular stress pathways related to dysregulation of glutathione metabolism, ferroptotic stress and ferroptosis have been implicated in failed repair of human AKI and pathogenesis in mouse models of AKI, (Fig. 5B), (18-20, 25, 28, 56). To investigate whether ferroptotic stress underlies the emergence and accumulation of damage-associated PT cells in addition to its known role in inducing cell death during maladaptive repair, we first tested the expression of the canonical anti-ferroptosis defense pathway, glutathione/GPX4 axis (Fig. 5B). In agreement with the underrepresentation of glutathione metabolic process in damage-associated PT cells, the genes encoding the glutathione/GPX4 defense pathway were markedly down-regulated in this PT cell state (DA-PT) compared to differentiated PT cells (PT), suggesting that damage-associated PT cells are potentially vulnerable to ferroptotic stress (Fig. 5C).

We then analyzed the expression of ferroptotic stress biomarkers such as malondialdehyde (MDA, a lipid peroxidation product) and acyl-CoA synthetase long-chain family member 4 (ACSL4), which also regulate cellular sensitivity to ferroptosis (21, 22, 56–59). A recent pharmacological inhibitor study showed that ACSL4 is a reliable maker for ferroptotic stress in murine model of ischemic AKI (24). We identified robust upregulation of *Acsl4* in damage-associated PT cells in dot-plots (Fig. 5C). The co-expression of markers for damage-associated PT cells and ferroptotic stress was confirmed by immunofluorescence for SOX9, MDA, and ACSL4 (Fig. 5, D-G). We found that severe ischemia (30 min) induces more expression of ferroptotic stress markers in SOX9*^+^* cells than mild ischemic injury (20 min) (Fig. 5, E and G). These data demonstrate that SOX9^+^ damage-associated PT cells undergo high ferroptotic stress after severe ischemic injury.

To address whether the emergence of damage-associated PT cells is specific to IRI injury or appears in other cases of acute kidney injury, we investigated the co-expression of SOX9 and VCAM1 in models of toxic renal injury (aristolochic acid nephropathy, AAN) and obstructive renal injury (unilateral ureteral obstruction, UUO), which lead to severe fibrosis. By immunofluorescence analyses of SOX9 and VCAM1 co-expression, we found the emergence of damage-associated PT cells in both models (Fig. 6, A and C). Furthermore, the SOX9-positive tubular epithelial cells in these models showed co-expression of ACSL4, suggesting that ferroptotic stress of damage-associated PT cells is a conserved response to kidney injury across various etiologies (Fig. 6, B and C).

**Figure 6.**
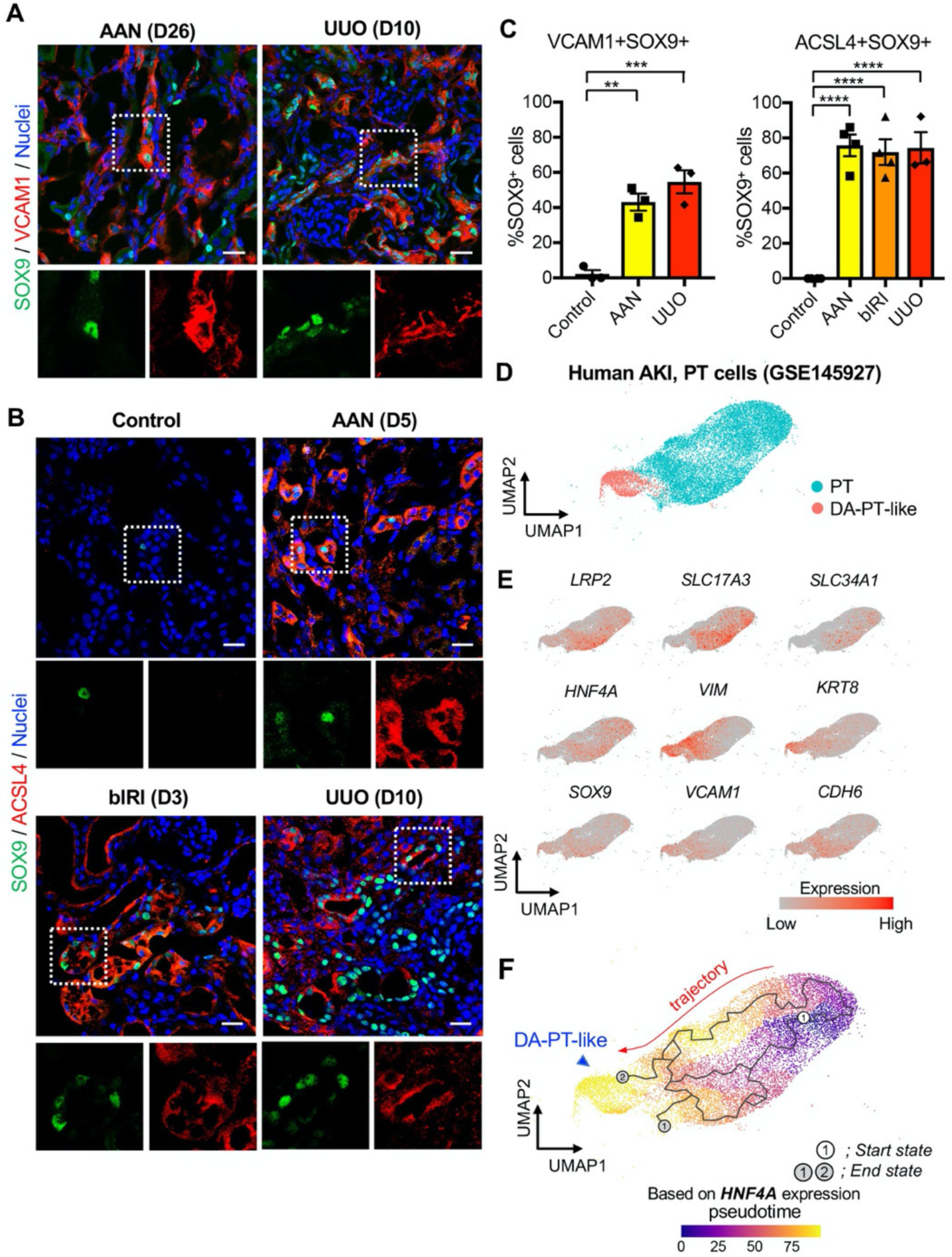
Damage-associated PT cells emerge after injury in mouse and human kidneys. **(A)** Immunostaining for SOX9 and VCAM1. Aristolochic acid nephropathy (AAN) and unilateral ureteral obstruction (UUO) models were used. Kidneys from wild type C57BL/6J mice were harvested on day 26 (D26) for AAN and day 10 (D10) for UUO. Insets: individual fluorescence channels of the dotted box area. (**B**) Immunostaining for SOX9 and ACSL4. bIRI, bilateral IRI model. Kidneys were harvested on day 3 (D3) for bIRI, day 5 (D5) for AAN, and day 10 (D10) for UUO. Insets: individual fluorescence channels of the dotted box area. (**C**) Quantification of double-positive cells in total SOX9^+^ cells from panel (A) and (B). Scale bars, 20 μm. N = 3-4. \*\**P* < 0.01; \*\*\**P* < 0.001; \*\*\*\**P* < 0.0001, one-way ANOVA with post hoc multiple comparisons test. (**D**) UMAP of the human proximal tubular cells from AKI kidneys. (**E**) UMAP plots showing the expression of indicated genes. Note that damage-associated PT cells (DA-PT-like) show increased gene expressions of markers for mouse damage-associated PT cells (*SOX9*, *VCAM1*, *CDH6*) and reduced expression of homeostatic genes (*LRP2*, *SLC17A3*, *SLC34A1*, and *HNF4A*). (**F**) Pseudotime trajectory analysis of PT clusters (PT and DA-PT-like cells). A region occupied with *HNF4A*^high^ cells were set as a starting state. Arrow, predicted trajectory from PT cells to DA-PT-like cells.

We then investigated whether molecularly similar damage-associated PT cells can be observed in human AKI. We analyzed scRNA-seq data from biopsy samples of two transplanted human kidneys with evidence of AKI and acute tubular injury but no evidence of rejection (GSE145927; Fig. 6D and Fig. S12A), (60). We found a cell population that is highly enriched for genes expressed in mouse damage-associated PT cells, including *SOX9*, *VCAM1*, and *CDH6*. This cellular population also showed decreased expressions of homeostatic PT genes (*SLC34A1*, *SLC17A3*, *LRP2,* and *HNF4a*) (Fig. 6, D, and E). Trajectory inference using Monocle 3 suggests that damage-associated PT cells emerge from mature differentiated PT cells that show high expressions of homeostatic genes in human kidneys (PT to DA-PT-like in Fig. 6F). Interestingly, the glutathione metabolic gene signature is high in mature PT cells and decreases along the trajectory to DA-PT-like cells (Fig. S12C). These data suggest that the emergence of damage-associated PT cells is a mechanism of acute kidney injury and repair that is shared by humans and mice.

### Genetic induction of ferroptotic stress results in accumulation of inflammatory PT cells after mild injury

Our data suggest that severe injury, which induces more oxidative and ferroptotic stress than mild injury, causes the accumulation of inflammatory damage-associated PT cells and worsens long-term renal outcomes. We hypothesized that ferroptotic stress plays a crucial role in driving the accumulation of inflammatory PT cells and promoting maladaptive repair in addition to triggering cell death (ferroptosis). To test this hypothesis, we generated a mouse model that selectively and conditionally deletes *Gpx4* in *Sox9*-lineage cells (*Sox9^IRES-CreERT2^*; *Gpx4^flox/flox^*, hereafter cKO; Fig. 7A). Genetic deletion of *Gpx4* robustly induces ferroptotic stress and triggers ferroptosis (20, 26). In this mouse line, exons 4-7 of the *Gpx4* allele, which include the catalytically active selenocysteine site of the GPX4 protein, is deleted in a tamoxifen-inducible manner selectively in *Sox9*-lineage cells. We subjected the cKO mice and littermate control mice to mild renal ischemic stress (ischemic time 22 min). This condition induces robust *Sox9-CreERT2* expression, but does not induce the failed renal repair phenotype in control mice (*Gpx4* ^flox/flox^). We induced *Gpx4* deletion at the time of injury by tamoxifen injection (Fig. 7A). The littermate control mice were subjected to the same renal ischemic stress and tamoxifen. We confirmed successful deletion of GPX4 protein by immunofluorescence (Fig. S13, B and C) and found that expression of the ferroptotic stress marker ACSL4 was increased on day 21 post-IRI in cKO kidneys compared to littermate kidneys that underwent the same ischemic stress (Fig. S13, D and E). Contralateral uninjured kidneys from cKO mice only showed a minimum deletion of GPX4 as the *Sox9-CreERT2* activity is not induced in non-injured kidneys (See Fig. 3B for CLK), (13).

**Figure 7.**
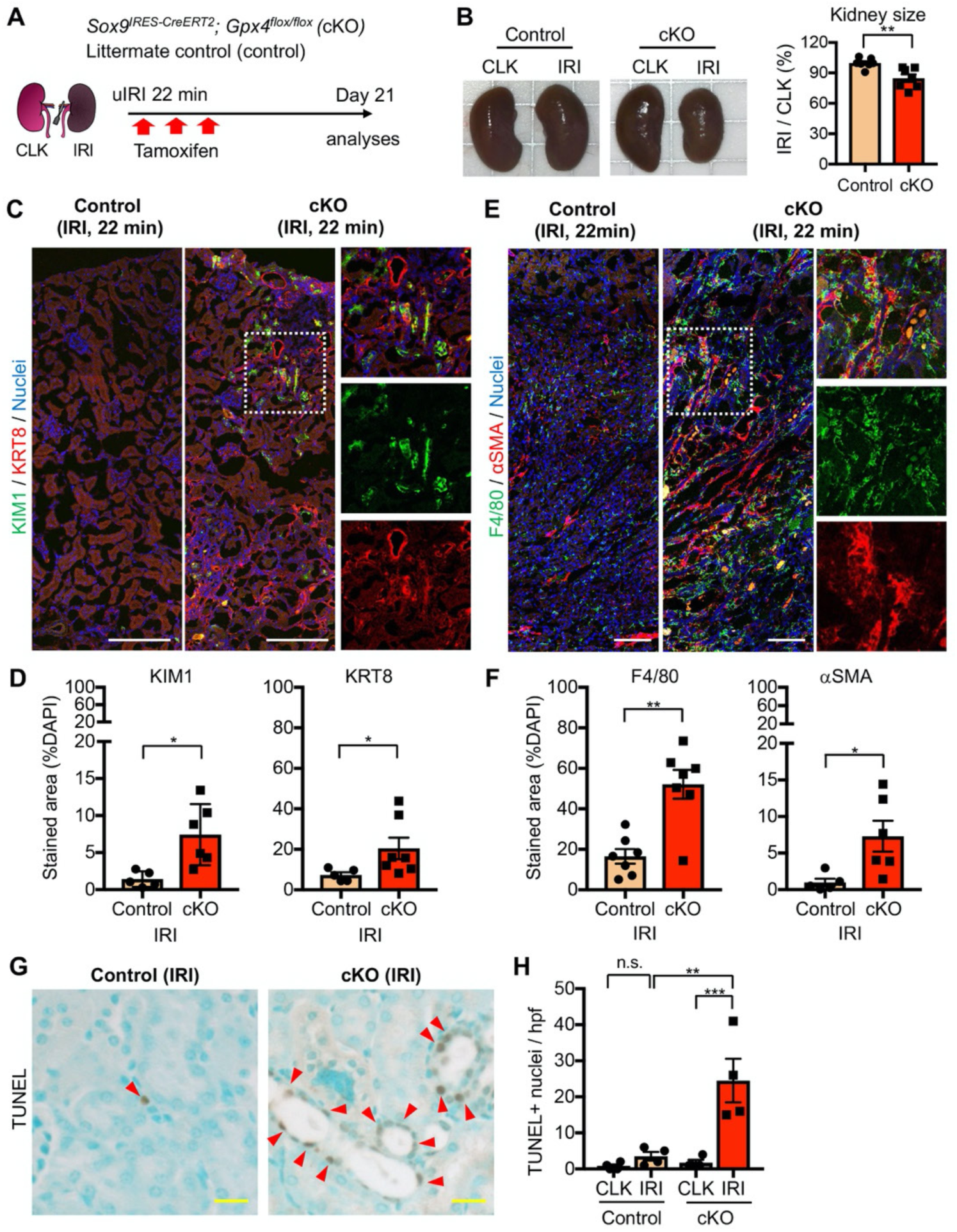
Genetic induction of ferroptotic stress to *Sox9*-lineage cells augments kidney injury. (**A**) Experimental workflow for *Gpx4* deletion in *Sox9*-lineage cells. uIRI, unilateral IRI (ischemic time 22 min). Kidneys were harvested on day 21 post-IRI. cKO mice and their littermate controls were subjected to the same ischemic stress and tamoxifen treatment. *Gpx4* is deleted in *Sox9-*lineage cells after IRI with tamoxifen administration. (**B**) The deletion of *Gpx4* results in renal atrophy. Relative size of post-IRI kidneys compared to contralateral kidneys (CLK) was quantified. Control, littermate control. N = 7. (**C**) and (**D**) Immunostaining for tubular injury markers (KIM1 and KRT8). IRI kidneys from cKO and control littermates are shown. CLK did not show KIM1 or KRT8 staining (See Fig. S15 for CLK data). Quantification of KIM1 or KRT8-positive area over the DAPI^+^ area is shown in (D). N = 5-7. (**E**) and (**F**) Immunostaining for F4/80 and *α*SMA. IRI kidneys from cKO and control littermates are shown. Quantification of F4/80 or *α*SMA-positive area over the DAPI^+^ area is shown in (F). N = 5-7. Insets: individual fluorescence channels of the dotted box area. (**G**) and (**H**) TUNEL staining for evaluating cell death. Quantification of TUNEL-positive nuclei is shown in (H). N = 4. Arrowheads, TUNEL^+^ nuclei. Unpaired t-test for (D) and (F). Abbrev: hpf, high power field. One-way ANOVA with post hoc multiple comparison test for (H). Scale bars, 100 μm in (C) and (E); 20 μm in (G).

The post-ischemic cKO kidneys were atrophic and showed severe tubular injury on histological evaluation on day 21 and exhibited marked accumulation of KIM1^+^KRT8^+^ injured tubular cells (Fig. 7, B-D and Fig. S14). By contrast, control littermate kidneys that underwent the same ischemic stress exhibited resolution of histological changes and fewer KIM1^+^KRT8^+^ cells (Fig. 7, C and D, and Fig. S14). Contralateral kidneys from both genotypes showed neither increased KIM1 nor KRT8 expression (Fig. S15A and S15C). The post-ischemic cKO kidneys also exhibited massive accumulation of F4/80^+^ macrophages, *α*SMA^+^ myofibroblasts, and increased collagen synthesis (Fig. 7, E-F; Fig. S15B and S15C). Then, we assessed the number of cell death by terminal deoxynucleotidyl transferase-mediated dUTP nick end labeling (TUNEL) assay, which detects ferroptotic cell death in *Gpx4*-deleted tissues (26). Consistent with the known role of GPX4 to prevent ferroptosis, genetic deletion of *Gpx4* led to the increased TUNEL^+^ tubular epithelial cells in cKO kidneys (Fig. 7, G and H; See Fig. S16 for CLK). Collectively, these data indicate that genetic induction of ferroptotic stress in *Sox9*-lineage cells is sufficient to prevent normal renal repair after mild ischemic injury and to mimic the failed renal repair phenotype observed after severe ischemic injury.

We then investigated if the number of damage-associated PT cells was increased in the *Gpx4* cKO kidneys after mild ischemic injury. While VCAM1 is strongly induced in damage-associated PT cells and serves as a reliable marker, it is also expressed weakly in F4/80^+^ macrophages and endomucin (EMCN)^+^ endothelial cells after kidney injury (Fig. S6A; See UMAP). For the precise quantification of damage-associated PT cells, we co-stained the kidneys with VCAM1, EMCN, and F4/80, and scored VCAM1^+^F4/80^−^EMCN^−^ cells as damage-associated PT cells (Fig. 8, B and C; and Fig. S15D). Supporting our hypothesis, we observed increased numbers of VCAM1^+^EMCN^−^F4/80^−^ cells in post-ischemic cKO kidneys on day 21, while the value was at a baseline level in control littermate kidneys that underwent the same mild ischemic stress (Fig. 8, B and C). We further employed a genetic fate-mapping strategy in *Gpx4*-deficient *Sox9*-lineage cells by generating a mouse line that harbors *Sox9^IRES-CreERT2^*; *Gpx4^flox/flox^; Rosa26^tdTomato^* alleles. Confocal imaging identified the colocalization of tdTomato (*Sox9*-lineage) and VCAM1 and ACSL4 in the post-IRI cKO kidneys (Fig. 8D). Other molecular markers of damage-associated PT cell state, such as *Cdh6* and *Sox9*, were also increased in cKO kidneys on day 21 post-IRI (Fig. 8E). These VCAM1^+^ cells were also positive for SOX9 (Fig. 8F). *In situ* hybridization confirmed robust *Cdh6* expression in tubular epithelial cells in post-ischemic cKO kidneys (Fig. 8G and Fig. S15E). These data substantiate our model that ferroptotic stress drives the accumulation of damage-associated PT cells by preventing redifferentiation of these transient inflammatory epithelial cells into normal PT cell state and augments renal inflammation and fibrosis (Fig. 8H).

**Figure 8.**
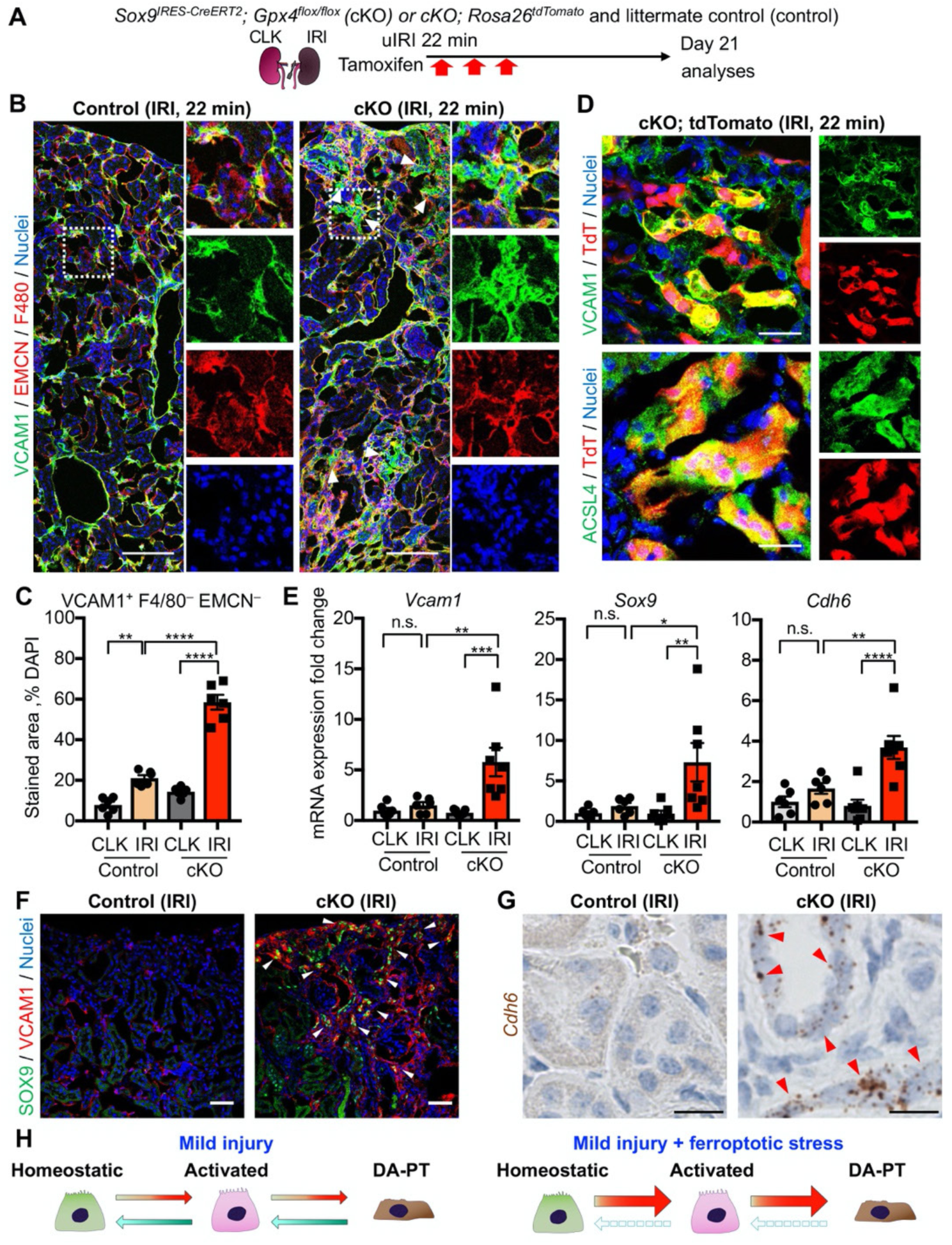
Genetic induction of ferroptotic stress induces the accumulation of damage-associated PT cells after mild injury. (**A**) Schematic representation of experimental workflow. tdTomato-lineage tracing was employed to detect *Sox9*-lineage cells. cKO mice and their littermate controls were subjected to the same ischemic stress (ischemic time, 22 min) and tamoxifen treatment. (**B**) and (**C**) Immunostaining for VCAM1, EMCN (endomucin), and F4/80. IRI kidneys from cKO and control littermates (control) are shown. Quantification of VCAM1^+^EMCN^−^F4/80^−^ area over the DAPI^+^ area is shown in (C). N= 6. See Fig. S15 for CLK data. (**D**) Immunostaining for VCAM1, ACSL4, and native tdTomato (TdT) fluorescence. Insets: individual fluorescence channels. (**E**) Real-time PCR analyses of indicated gene expression. Whole kidney lysates were used. N= 6-7. \**P* < 0.05; \*\**P* < 0.01; \*\*\**P* < 0.001; \*\*\*\**P* < 0.0001, one-way ANOVA with post hoc multiple comparisons test for (C) and (E). (**F**) Immunostaining for SOX9 and VCAM1. Arrowheads indicate double-positive cells (damage-associated PT cells). (**G**) ISH for *Cdh6* expression. Red arrowheads indicate *Cdh6*-positive renal tubular cells. Scale bars, 100 μm in (B); 20 μm in (D); 50 μm in (F); and 10 μm in (G). See Fig. S15 for CLK data. (**H**) Schematic illustration of PT cell state dynamics. Differentiated/mature PT cells are activated, transit into a damage-associated inflammatory PT cell state (DA-PT), and redifferentiate to their original state after mild injury. Ferroptotic stress prevents the redifferentiation of damage-associated PT cells into normal PT cell state, leading to the accumulation of the pathologic PT cells that actively produce inflammatory signals.

## Discussion

By using complementary scRNA-seq and mouse genetic approaches in several experimental models of renal injury and repair, our study revealed novel mechanisms regulating proximal tubular cell states that underlie renal repair and regeneration. By detailed characterization of damage-associated PT cells in our single-cell map of failed repair, we identified that this PT state significantly downregulates the canonical anti-ferroptosis defense pathway, making them potentially vulnerable to ferroptotic stress. Genetic induction of ferroptotic stress after mild injury was sufficient to prevent the redifferentiation of damage-associated PT cells into the normal PT cell state, leading to the accumulation and persistence of inflammatory PT cells that promote maladaptive repair. Our data collectively advances our understanding of the ferroptotic cell death pathway by identifying a novel role of ferroptotic stress in promoting and accumulating pathologic cellular state beyond its known role to trigger non-apoptotic regulated cell death (ferroptosis).

Unbiased clustering of cells clearly separates damage-associated PT cells from homeostatic and activated differentiated PT cells, indicating that damage-associated PT cells represent a unique cellular status. We also found a molecularly similar PT cell state in kidneys of patients with acute kidney injury. Similar to our current findings, we and others have identified the emergence of molecularly distinct epithelial cells during the process of lung injury and repair (61–63). These novel transient cells are termed as pre-alveolar type-1 transitional cell state (PATS), alveolar differentiation intermediate, and damage-associated transient progenitors. They originate from alveolar type 2 epithelial cells and differentiate into type 1 alveolar epithelial cells (61–63). PATS and PATS-like cells in humans accumulate during failed lung repair and fibrosis (61), as in the case of maladaptive repair of kidneys. Molecular mechanisms underlying the accumulation of these transitional cell state include hypoxia, inflammation, and DNA damage. All of these pathways promote maladaptive renal repair by altering PT cell states (3, 5, 7, 54). These data suggest that the emergence of molecularly distinct epithelial cell states and their persistence/accumulation is a general mechanism of maladaptive repair in multiple organs across mice and humans.

The complexity of proximal tubular cell states in renal injury and repair processes has been recently identified at single-cell resolution (10, 11). A recent study investigated PT cellular heterogeneity using single-nucleus RNA sequencing in a mouse model of bilateral renal IRI. The paper revealed multiple novel PT cellular states, ranging from severely injured cells, cells repairing from injury, and cells undergoing failed repair (10). Interestingly, the damage-associated PT cells reported here shares some of the transcriptional signatures with so-called failed repair proximal tubular cells (FR-PTC), such as *Vcam1, Cp, Akap12, and Dcdc2a* among the Top 20 transcriptional signature of FR-PTC. In contrast, we also found some differences between the damage-associated PT cells and FR-PTC. The most highly expressed genes in FR-PTC (ex. *Kcnip4, Dock10, Pdgfd, Erbb4, and Psd3*) were not expressed in damage-associated PT cells and *vice versa*. Moreover, damage-associated PT cells act like a transient cell state. They redifferentiate to the homeostatic PT cell state after mild injury while they accumulate after severe injury. Damage-associated PT cells may represent a broad transient cell state, including FR-PTC.

Another study profiled juvenile (4 week-old) mouse kidneys that underwent 30-min unilateral IRI (11). Unlike adult kidneys, the kidneys at this stage showed marked regenerative ability and showed successful repair. The study found transient induction of nephrogenic transcriptional signature (ex. *Sox4*, *Cd24a*, *Npnt*, *Lhx1, Osr2, Foxc1, Hes1, Pou3f3, and Sox9*) in damaged PT cells during the injury–repair process (11). While our transcriptional analyses of damage-associated PT cells indicate they are in a dedifferentiated state, they do not show a nephrogenic signature at the level of damaged juvenile kidneys (*i.e.,* damage-associated PT cells were positive for *Sox4, Npnt*, and *Cd24a*, and *Sox9*, but negative for *Lhx1, Osr2, Foxc1, Hes1, and Pou3f3*). The differences in reactivation of developmental genes between adult and juvenile kidneys may underlie the age-dependent decline of reparative capacity of mouse and human kidneys. Future studies testing the proximal tubular heterogeneity and spatiotemporal dynamics in additional renal injury models in young and aged animals may offer further insights into molecular mechanisms governing proximal tubular cell plasticity and identify therapeutic targets.

It has been largely believed that ferroptotic stress reduces functional renal epithelial cells by intercellular propagation of ferroptotic cell death (synchronized cell death) and induces so-called necroinflammation (25, 26, 59, 63). Consistent with this notion, we observed the accumulation of TUNEL^+^ tubular epithelial cells in cKO kidneys. In addition to inducing ferroptosis in some tubular epithelial cells, to our surprise, our genetic knockout studies showed that excess ferroptotic stress in regenerating PT cells drives the accumulation, but not reduction, of damage-associated PT cells that augment renal inflammation. Specific gene-expression signatures indicate that damage-associated PT cells are not merely severely injured cells on the pathway to cell death but a unique functional cell state. The cells are enriched for expression of renal developmental genes such as *Sox4* and *Sox9*. SOX9 is a previously described transcription factor essential for renal repair (12, 13), and SOX4 regulates epithelial mesenchymal transition in different disease contexts (64). Moreover, damage-associated PT cells are actively involved in renal inflammation by interacting with myeloid cells through producing cytokines and chemokines. Thus, ferroptotic stress not only promotes the alteration of cell state but makes it irreversible, leading to the pathologic accumulation of cells that actively produce inflammatory and fibrogenic signals.

In summary, our study broadens the roles of ferroptotic stress from one that is restricted to the induction of regulated cell death (ferroptosis) to include the promotion and accumulation of pathologic cell state that underlie maladaptive repair. Understanding the molecular mechanisms by which ferroptotic stress controls these processes *in vivo* would open a new avenue for currently available and prospective anti-ferroptotic reagents to enhance tissue repair/regeneration in multiple organs. Our studies provide a scientific foundation for future mechanistic and translational studies to enhance renal repair and regeneration by modulating anti-ferroptotic stress pathways to prevent AKI to CKD transition in patients.

## Materials and Methods

### Animals

All animal experiments were approved by the Institutional Animal Care and Use Committee at Duke University and performed according to the IACUC-approved protocol (A051-18-02 and A014-21-01) and adhered to the NIH Guide for the Care and Use of Laboratory Animals. The following mouse lines were used for our study; *Sox9^IRES-CreERT2,^* (46), *Rosa26^tdTomato^* (Jackson lab, stock #007914), (65), *Gpx4^flox^* (Jackson lab, stock# 027964), (66), and C57BL/6J (Jackson lab, stock #000664). Mice were backcrossed into a C57BL/6J background at least 3 times and maintained in our specific-pathogen-free facility. Timed deletion of the *Gpx4* gene and fate-mapping was achieved using *Sox9^IRES-CreERT2^* knock-in mouse line with 3 doses of intraperitoneal injections of tamoxifen (100 mg/kg body weight, Sigma, St. Louis MO) on alternate days. The first dose of tamoxifen was administered immediately before the surgical intervention. All tested animals were included in data analyses, and outliers were not excluded. To avoid confounding effects of age and strain background, littermate controls were used for all phenotype analyses of genetically modified mouse lines. Animals were allocated randomly into the experimental groups and analyses. The operators were blinded to mouse genotypes when inducing surgical injury models. To determine experimental sample sizes to observe significant differences reproducibly, data from our previous studies were used to estimate the required numbers. The number of biological replicates is represented by N in each figure legend. Experiments were performed on at least three biological replicates.

### Mouse models of renal injury and repair

Adult male mice aged between 8–16 weeks were used for all the models described below. The mice were euthanized, and kidneys were harvested for analyses. For the unilateral IRI (uIRI) model, ischemia was induced by the retroperitoneal approach on the left kidney for 20 min (mild IRI), 22 min (mild IRI in cKO studies), or 30 min (severe IRI) by an atraumatic vascular clip (Roboz, RS-5435, Gaithersburg, MD), as previously reported (27, 67). Mice were anesthetized with isoflurane and provided preemptive analgesics (buprenorphine SR). The body temperature of mice was monitored and maintained on a heat-controlled surgical pad. For the bilateral IRI (bIRI) model, ischemia was induced by the retroperitoneal approach on both kidneys for 22 min. The mice were received intraperitoneal injections of 500 μl of normal saline at the end of surgery. For the unilateral ureteral obstruction (UUO) model, the left ureter was tied at the level of the lower pole of the kidney, and the kidneys were harvested on day 10. For the aristolochic acid nephropathy (AAN) model, we used acute and chronic models, as we previously described (68). For the acute AAN model, 3 doses of 6 mg/kg body weight aristolochic acid in phosphate-buffered saline (PBS) were administered daily intraperitoneally to the male mice. For the chronic AAN model, 6 doses of 6 mg/kg body weight aristolochic acid in phosphate-buffered saline (PBS) were administered on alternate days over two weeks intraperitoneally to the male mice. The same volume of PBS was injected to control animals (68, 69). Contralateral kidneys (CLK), sham-treated kidneys, and vehicle-injected kidneys were used as controls depending on the models used. The numbers and dates of treatment are indicated in the individual figure legends and experimental schemes. Operators were blinded to mouse genotypes when inducing surgical injury models.

### Droplet-based scRNA-seq

Mice were transcardially perfused with ice-cold PBS, and the kidneys were harvested. The kidneys were dissociated with liberase TM (0.3 mg/mL, Roche, Basel, Switzerland, #291963), hyaluronidase (10 μg/mL, Sigma, H4272), DNaseI (20 μg/mL) at 37°C for 40 min, followed by incubation with 0.25% trypsin EDTA at 37°C for 30 min. Trypsin was inactivated using 10% fetal bovine serum in PBS. Cells were then resuspended in PBS supplemented with 0.01% bovine serum albumin. Our protocol yielded high cell viability (>95%) and very few doublets, enabling us to avoid the use of flow cytometry-based cell sorting. After filtration through a 40 μm strainer, cells at a concentration of 100 cells/μl were run through microfluidic channels along with mRNA capture beads and droplet-generating oil, as previously described (61, 70). cDNA libraries were generated and sequenced using HiSeq X Ten with 150-bp paired-end sequencing. Each condition contains the cells from three mice to minimize potential biological and technical variability. Detailed protocols for the computational analyses of scRNA-seq data are available in the supplementary method.

### Tissue Collection and Histology

Kidneys were prepared as described previously (27, 51). For cryosections (7 μm), the tissues were fixed with 4% paraformaldehyde in PBS at 4°C for 4 hours and then processed through a sucrose gradient. Kidneys were embedded in OCT compound for sectioning. For paraffin sections (5 μm), the tissues were fixed with 10% neutral buffered formalin overnight at 4°C and processed at Substrate Services Core & Research Support at Duke. Sections were blocked (animal-free blocker with 0.5% triton x-100) for 30 min and incubated with the primary antibodies overnight at 4°C. Primary antibodies used were as follows: SOX9 (Abcam, Cambridge, UK, ab196450 or ab185966, 1:200), KIM1 (R&D Systems, Minneapolis, MN, AF1817, 1:400), NGAL (Abcam, ab70287, 1:400), F4/80 (Bio-rad, Hercules, CA, MCA497G, 1:200), *α*-SMA (Sigma, C6198, 1:200), LTL (Vector, Burlingame, CA, B-1325 or FL-1321, 1:200), KRT8 (DSHB, TROMA-I, 1:200), MDA (Abcam, ab6463, 1:200), ACSL4 (Abcam, ab204380 or ab155282, 1:200), EMN (Abcam, 106100, 1:200), VCAM1 (CST, 39036S or 33901S, 1:100), and GPX4 (Abcam, ab125066, 1:200). Alexa Fluor-labeled secondary antibodies were used appropriately. Nuclei were stained with DAPI (1:400, Sigma). Heat-induced antigen retrieval was performed using pH 6.0 sodium citrate solution (eBioscience). Experiments for RNAScope *in situ* hybridization (Advanced Cell Diagnostics, ACD, Newark, CA) was performed as recommended by the manufacturer. Mm-Cdh6 (ACD, 519541) was used. Images were captured using Axio imager and 780 confocal microscopes (Zeiss, Oberkochen, Germany). Paraffin-sections were stained with hematoxylin and eosin (H&E). The kidney injury score was calculated as we previously reported (68). TUNEL staining was performed following the manufacturer’s instruction (Abcam, ab206386). To ensure the TUNEL signal’s specificity, we used sections treated with DNase I as a positive control and a section treated without terminal deoxynucleotidyl transferase as a negative control, as recommended by the manufacturer. Sections were counterstained with methyl green. More than three randomly selected areas from at least three kidneys were imaged and quantified using ImageJ (51). The stitched large area was used for quantification to alleviate the selection bias in the acquisition of images. All representative images were from more than 3 kidneys tested.

### Statistical analysis

Statistical analyses were conducted using GraphPad Prism software. Two-tailed unpaired Student’s t-test was used for two groups, and one-way analysis of variance (ANOVA) followed by Sidak multiple comparison test was used for more than two groups. All results are represented as means ± SE. A *P* value less than 0.05 was considered statistically significant.

Additional protocols are available in the supplementary method.

## Acknowledgments

We thank Drs. Brigid Hogan and Myles Wolf for critical advice and helpful suggestions on the manuscript. We also thank Dr. Helene F. Kirshner (Duke Center for Genomic and Computational Biology) for her bioinformatical support on our single-cell transcriptome dataset. The monoclonal antibody against keratin 8 (TROMA-1, developed by Drs. P. Brulet and R. Kemeler) was obtained from the Developmental Studies Hybridoma Bank, created by the NICHD of the NIH and maintained at the Department of Biology, The University of Iowa. We thank Drs. Tetsuhiro Yokonishi (Duke University), Leslie Gewin and Kensei Taguchi (Vanderbilt University), and members of the Crowley lab for their technical advice. This study was supported by grants from the National Institute of Diabetes and Digestive and Kidney Diseases (R01 DK123097), the American Society of Nephrology Carl W. Gottschalk Career Developmental Grant, and Duke Nephrology Start-up Fund to TS. SI, YK, and KI are supported in part by fellowship grants from the American Heart Association, Japan Society for the Promotion of Science, and the Astellas Foundation for Research on Metabolic Disorders, respectively. Imaging was performed at the Duke Light Microscopy Core Facility supported by the shared instrumentation grant (1S10RR027528-01).

## Data availability

Single-cell RNA-seq data of this study have been deposited in the Gene Expression Omnibus (GEO; GSE161201). Primer sequence information is available in the supplementary file.

## Author contributions

Y.K., S.I., T.S., and P.R.T designed the single-cell transcriptomics experiments and analyzed the data; S.I., Y.K., K.I., S.A.S., S. H. and T.S. performed experiments; L.L.O, S.D. C., L.B., A. T., P.R. T contributed new protocols/reagents/analytical tools; S.I., Y.K., K.I., and T.S. analyzed the data; T.S., S.I., Y.K., S.D.C., A.T., and P.R.T wrote the manuscript. T. S. conceived and supervised the study. All authors read, commented, and approved the manuscript.

## Supplementary Methods

### Computational analyses of scRNA-seq data

#### 1) Data preprocessing, unsupervised clustering, and cell type annotation of Drop-Seq data

Analysis of the scRNA-seq of mouse kidneys was performed by processing FASTQ files using dropSeqPipe v0.3 and mapped on the GRCm38 genome reference with annotation version 91. Unique molecular identifier (UMI) counts were then further analyzed using an R package Seurat v.3.06 for quality control, dimensionality reduction, and cell clustering (1). The scRNA-seq matrices were filtered by custom cutoff (genes expressed in >3 cells and cells expressing more than 500 and less than 3000 detected genes were included) to remove potential empty droplets and doublets. Relationships between the number of UMI/cell and genes/cell were comparable across the condition (Fig. S3A). After quality control filtration and normalization using SCTransform (2), UMI count matrices from post-IRI kidneys and homeostatic kidneys were integrated using Seurat’s integration and label transfer method, which corrects potential batch effects (1). The integrated dataset was used for all the analyses. To remove an additional confounding source of variation, the mitochondrial mapping percentage was regressed out. The number of principal components (PC) for downstream analyses were determined using elbow plot to identify knee point, and we included the first 25 PCs for the downstream analyses. A graph-based clustering approach in Seurat was used to cluster the cells in our integrated dataset. The resolution was set at 1.0 for the mouse integrated dataset. Cluster-defining markers for each cluster were obtained using the Seurat FindAllMarkers command (genes at least expressed in 25% of cells within the cluster, log fold change> 0.25) with the Wilcoxon Rank Sum test (Table S1). Based on the marker genes and manual curation of the gene expression pattern of canonical marker genes in UMAP plots (Fig. S4), we assigned a cell identity to each cluster. Ambiguous clusters were shown as unknown. We manually combined 3 clusters of differentiated proximal tubular cells (PT, S1/S2 and PT, S2/S3; Fig. S4) into 1 cluster (PT) to generate a more coarse-grained cell-type annotation and data visualization. We also combined 3 clusters of endothelial cells (Endo-1, Endo-2, and Endo-3; Fig. S4) into one cluster (Endo) for data visualization.

#### 2) Data preprocessing, unsupervised clustering, and cell type annotation of mouse neonatal kidneys

The RDS files for mouse neonatal kidneys (postnatal day 1) were obtained from Gene Expression Omnibus (GEO accession number: GSE94333, GSM2473317), (3). Data were analyzed as in our mouse kidney dataset using Seurat and SCTransform (1, 2). We included the first 17 PCs for the downstream analyses of mouse neonatal kidneys. A graph-based clustering approach in Seurat was used to cluster the cells. The resolution was set at 0.8. Based on the marker genes and manual curation of the gene expression pattern of canonical marker genes in UMAP plots (Fig. S7), we assigned a cell identity to each cluster. The anchor genes for assigning cell identity were obtained from previous single-cell transcriptome analyses of the developing mouse kidneys (3, 4).

#### 3) Data preprocessing, unsupervised clustering, and cell type annotation of human kidneys

The RDS files for human kidneys were obtained from Gene Expression Omnibus (GEO accession number: GSE131882 and GSE145927), (5, 6). Normal human kidney data was originated from two macroscopically normal nephrectomy samples without renal mass (GSE131882; GSM3823939 and GSM3823941), (5). Human AKI kidney data was originated from two biopsy-samples of transplant kidneys with evidence of AKI and acute tubular injury but no evidence of rejection (GSE145927; GSM4339775 and GSM4339778), (6). Data were integrated and analyzed as in the mouse kidney analyses using Seurat’s integration method and SCTransform (1, 2). We included the first 25 PCs for the downstream analyses of human normal and AKI kidneys. A graph-based clustering approach in Seurat was used to cluster the cells. The resolution was set at 0.5 for normal human kidneys and 1.0 for the human AKI kidneys. Based on the marker genes and manual curation of the gene expression pattern of canonical marker genes in UMAP plots (Fig. S11 and S12), we assigned a cell identity to each cluster. The anchor genes for assigning cell identity were obtained from previous single-cell transcriptome analyses of the human kidneys (5–7).

#### 4) Differential gene expression analyses and Gene Ontology (GO) enrichment analyses

To predict the cellular functions based on enriched gene signature, we performed gene-ontology enrichment analyses. Differentially expressed genes obtained using FindMarkers command in Seurat were used for identifying signaling pathways and gene ontology through Enricher (Table S2 and S3; Figure S5B), (8). To visualize the overrepresented signaling pathways, scaled data in the integrated Seurat object were extracted. Then, mean values of the scaled score of gene members in each GO class were calculated and shown in UMAP (9). The gene member lists of signaling pathways were obtained from AmiGO 2 (10). Log_2_ fold changes and *P*-values of each gene extracted using FindMarkers command in Seurat with Wilcoxon rank sum test were shown in a volcano plot using an R package EnhancedVolcano v1.4.0 (https://github.com/kevinblighe/EnhancedVolcano), (Fig. S5B).

Top 100 genes in mature and early PT cell clusters were obtained using the “FindMarkers” command in Seurat. These genes were visualized on the UMAP plots using the scaled score as in GO class visualization.

#### 5) RNA velocity analyses

To infer future states of individual cells, we performed RNA velocity analyses(11) using single-time point dataset of post-IRI kidney on day 7. The aligned BAM files were used as input for Velocyto to obtain the counts of unspliced and spliced reads in loom format. Cell barcodes for the clusters of interests (PT and DA-PT) were extracted and utilized for velocyto run command in velocyto.py v0.17.15, as well as for generating RNA velocity plots using velocyto.R v0.6 in combination with an R package SeuratWrappers v0.2.0 (https://github.com/satijalab/seurat-wrappers). Twenty-five nearest neighbors in slope calculation smoothing were used for RunVelocity command.

#### 6) Pseudotime trajectory analyses

To infer the dynamic cellular process during injury and repair, we performed single-cell trajectory analyses. We first extracted the clusters of interests (PT and DA-PT) from our integrated Seurat object of mouse kidneys and utilized for Monocle 3 (version 0.2.3.0) analyses with default parameters to identify a pseudotime trajectory with SeuratWrappers v0.2.0 (12, 13). We set the starting states in two different approaches. We used the UMAP space area occupied by cells from the earliest time point of IRI kidneys (6 hr post-IRI, Fig. 1F) and the area occupied by the cells with high expression of genes that are highly expressed in differentiated PT cells, such as *Slc34a1* (Fig. S8B) as the starting state, respectively. Both approaches resulted in similar trajectory inference. For the human AKI dataset, we extracted the clusters of interests (PT and DA-PT-like) from our integrated Seurat object and applied the Monocle 3 algorithm with default parameters. We used the UMAP space area occupied by the cells with high expression of homeostatic genes (*HNF4A*), (Fig. 6F).

#### 7) Intercellular communication analyses using NicheNet

To predict the intercellular communication process between damage-associated PT (DA-PT) cells and myeloid cells (monocytes and macrophages), we performed NicheNet analyses based on the analytical pipeline (https://github.com/saeyslab/nichenetr/blob/master/vignettes/seurat_wrapper.md) using an R package nichenetr (version 1.0.0) with default parameters (14). Based on high enrichment of chemokines and cytokines in DA-PT cells and the observed positive association between the numbers of macrophages and DA-PT cells in severely injured kidneys, we surmised that they have a close molecular interaction. We used NicheNet to predict the ligand-receptor pairs that are most likely to explain the target gene expression in renal myeloid cells after IRI. We defined DA-PT cells as the “sender/niche” cell population and myeloid cells as the “receiver/target” cell population in our integrated Seurat object for these analyses. We defined the differentially expressed genes in monocytes or macrophages in IRI-kidneys compared to homeostatic kidneys as the gene sets of interest that were affected by predicted ligand-receptor interactions.

### RNA extraction and real-time quantitative PCR

Total RNA was extracted from kidneys using the TRIzol reagent (Invitrogen, 15596026). 3 μg of total RNA was then reverse transcribed with Maxima H minus cDNA synthesis master mix (Invitrogen, M1662). Equivalent amounts of diluted cDNA from each samples were analyzed with Real-time PCR with the primers listed below using the Powerup SYBR Green reagent (Invitrogen, A25776) on a QuantStudio 3 real-time PCR systems (Thermo). 18S rRNA expression was used to normalize samples using the *ΔΔ*CT-method.

*Sox9*: Fw-GAGCCGGATCTGAAGAGGGA, Rv-GCTTGACGTGTGGCTTGTTC

*Vcam1*: Fw-TCTTACCTGTGCGCTGTGAC, Rv-ACTGGATCTTCAGGGAATGAGT

*Cdh6*: Fw-CCAATATTCACCAAGGACGTTTA, Rv-CGTGACTTGGACCACAAATG

*Acsm2*: Fw-CCAAGATGGCAGAACACTCC, Rv-TCAGAAGTACTCAGGCCTGTCC

*Icam1*: Fw-GCTACCATCACCGTGTATTCG, Rv-AGGTCCTTGCCTACTTGCTG

*Pdgfb*: Fw-CGAGGGAGGAGGAGCCTA, Rv-GTCTTGCACTCGGCGATTA

*Apoe*: Fw-TTGGTCACATTGCTGACAGG, Rv-AGCGCAGGTAATCCCAGAA

*Havcr1*: Fw-AAACCAGAGATTCCCACACG, Rv-GTCGTGGGTCTTCCTGTAGC

*Lcn2*: Fw-CAAGCAATACTTCAAAATTACCCTGTA, Rv-GCAAAGCGGGTGAAACGTT

*Acta2*: Fw-CCCACCCAGAGTGGAGAA, Rv-ACATAGCTGGAGCAGCGTCT

*Slc34a1*: Fw-CTCATTCGGATTTGGTGTCA, Rv-GGCCTCTACCCTGGACATAGA

*Krt8*: Fw-CTGAGCTTGGCAACATGC, Rv-ACGCTTGTTGATCTCATCCTC

*18S rRNA*: Fw-CGGCTACCACATCCAAGGAA, Rv-GCTGGAATTACCGCGGCT

### Mice and breeding

Genetically modified mouse lines were genotyped by PCR, using the following primers.

*Cre*, Fw: GTGCAAGTTGAATAACCGGAAATGG,

Cre, Rv: AGAGTCATCCTTAGCGCCGTAAATCAAT

*Gpx4 ^flox^*, wt: CTGCAACAGCTCCGAGTTC

*Gpx4 ^flox^*, common: CGGTGCCAAAGAAAGAAAGT

*Gpx4 ^flox^*, mut: CCAGTAAGCAGTGGGTTCTC

*Rosa26^tdTomato^*, Fw: CTGTTCCTGTACGGCATGG

*Rosa26^tdTomato^*, Rv-GGCATTAAAGCAGCGTATCC

*Rosa26^wt^*, Fw: AAGGGAGCTGCAGTGGAGTA

*Rosa26^wt^*, Rv: CCGAAAATCTGTGGGAAGTC.

---

**Fig. S1.**
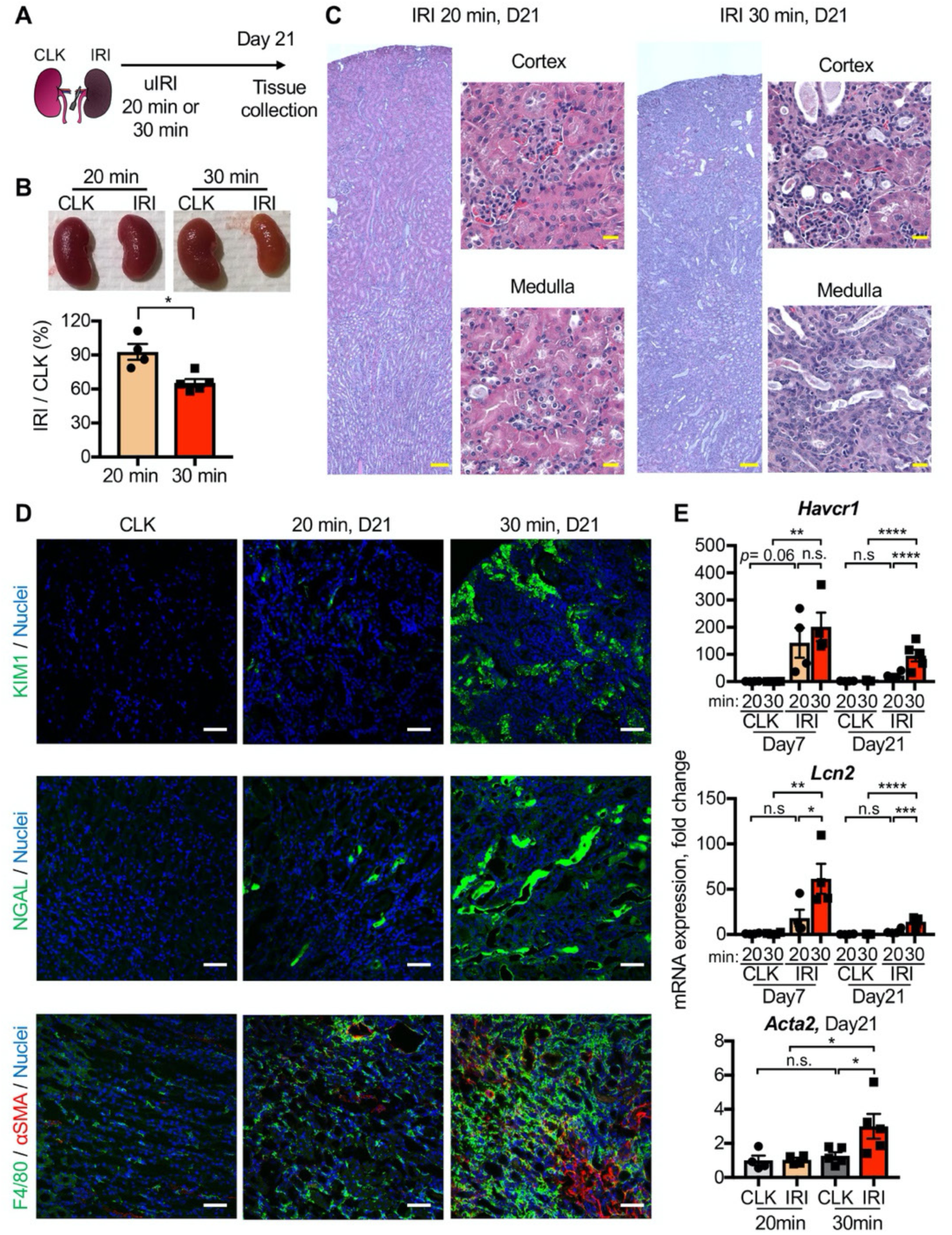
Characterization of severe and mild unilateral IRI models. (**A**) Experimental workflow for the mild and severe IRI models. Left kidneys from wild-type C57BL/6J mice were subjected to mild (20 min) and severe (30 min) ischemia. Contralateral kidneys (CLK) were used as controls. uIRI, unilateral IRI. (**B**) Severe IRI results in renal atrophy. Relative size of post-IRI kidney compared to contralateral kidney (CLK) was quantified. N = 4-5. (**C**) Hematoxylin-eosin staining of IRI kidneys on day 21 (D21). Note that severe-IRI resulted in tubular dilatation, flattening of tubular epithelial cells, cast formation, and inflammatory cell infiltration. (**D**) Immunofluorescence for KIM1, NGAL, F4/80, and *α*SMA. Severe IRI resulted in persistent expression of proximal and distal tubular injury markers. Kidney injury molecule 1, KIM1 (encoded by *Havcr1*) is a marker for proximal tubular injury. Neutrophil gelatinase-associated lipocalin, NGAL (encoded by *Lcn2*) is a marker for distal tubular injury. Note that post-severe IRI kidneys exhibit accumulation of F4/80^+^ cells (mostly macrophages) and *α*SMA^+^ myofibroblasts in renal interstitium, both are classical features of failed renal repair. (**E**) Real-time PCR analyses of indicated gene expression. *Acta2* gene encodes *α*SMA. N = 4-5. Scale bars: 100 μm in stitched images of (C); 20 μm in higher magnifications of (C); 50 μm in (D). \**P* < 0.05; \*\**P* < 0.01; \*\*\**P* < 0.001; \*\*\*\**P* < 0.0001. Unpaired Student’s t-test for (B) and one-way ANOVA with post hoc multiple comparisons test for (E).

**Fig. S2.**
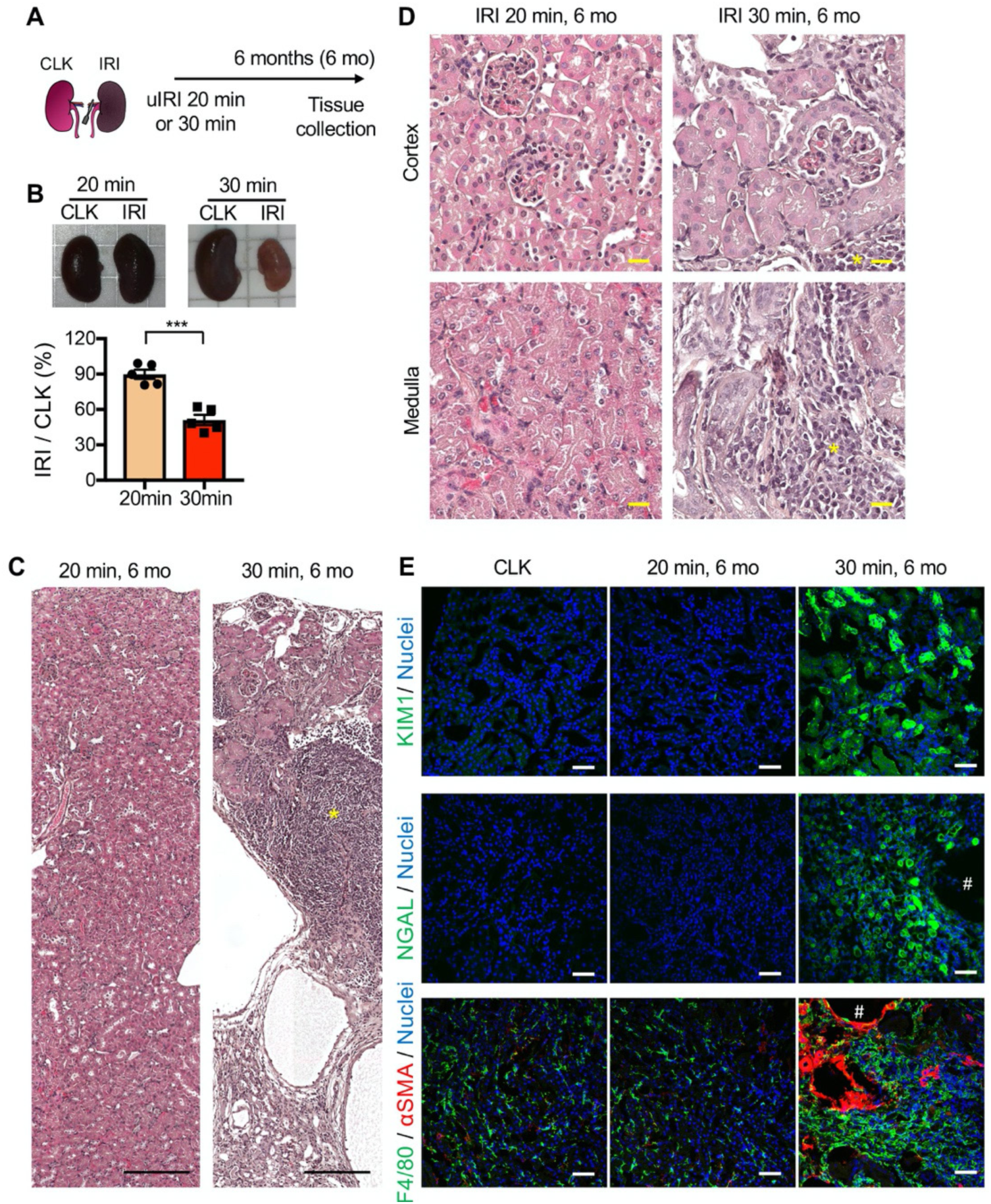
Severe IRI leads to cystic and atrophic kidneys 6 months after severe IRI. (**A**) Experimental workflow for mild and severe IRI models with long-term observation. (**B**) Severe IRI (30 min ischemia) results in marked renal atrophy on 6 months post-IRI. Relative size of post-IRI kidney compared to contralateral kidney (CLK) was quantified. N = 5. Unpaired Student’s t-test. (**C**) and (**D**) Hematoxylin-eosin staining of IRI kidneys. Note that severe-IRI resulted in flattened and necrotic tubular epithelial cells with massive infiltration of inflammatory cells, which occupied the renal parenchyma. * indicates clusters of inflammatory cells, occupying renal parenchyma. (**E**) Immunofluorescence for KIM1, NGAL, F4/80, and *α*SMA. Severe IRI resulted in persistent expression of proximal and distal tubular injury markers (KIM1 and NGAL, respectively). Note that post-severe IRI kidneys exhibit accumulation of F4/80^+^ myeloid cells (mostly macrophages) and *α*SMA^+^ myofibroblasts in renal interstitium. Small cystic structures (#) were encircled by myofibroblasts. Scale bars: 20 μm in (C), 200 μm in (D) and 50 μm in (E).

**Fig. S3.**
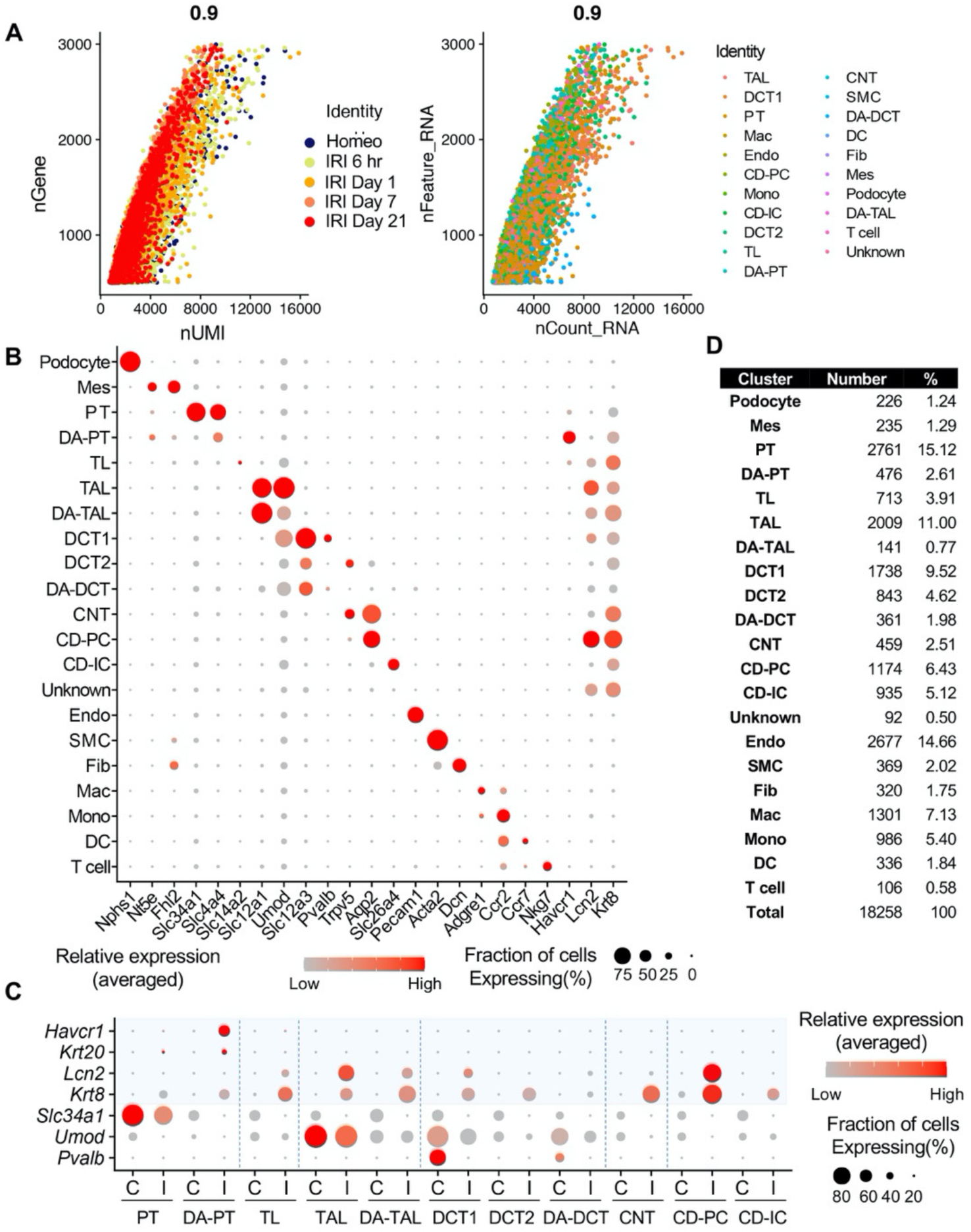
scRNA-seq identifies major cell types in homeostatic and post-IRI kidneys. (**A**) Pearson correlation plot showing the linear relationship between the number of genes (nGene) and unique molecular identifiers (nUMI). Experimental conditions and cell types are color-coded. (**B**) Dot plot shows the gene expression patterns of cluster-enriched canonical markers. Note that damage-associated tubular epithelial cells (DA-PT, DA-TAL, and DA-DCT) have reduced expression of canonical cellular markers. (**C**) Tubular injury marker gene expressions are selectively observed in damaged kidneys (IRI) but not in homeostatic control kidneys. “C” indicates cells from the control homeostatic kidneys. “I” indicates cells from IRI kidneys from all time points. Proximal tubule-specific injury markers (*Havcr1*, *Krt20*), distal tubule-specific injury markers (*Lcn2*), and pan-tubular injury marker (*Krt8*) are shown. Both homeostatic and activated cells show high canonical cell type marker gene expressions, such as *Slc34a1* in PT, but damage-associated clusters (DA-PT, DA-TAL, DA-DCT) show reduced homeostatic gene expressions and increased damage-induced gene expressions (See Fig. 1A). (**D**) Our single-cell preparation resulted in high yields of podocytes (1.24%) and endothelial cells (14.66%). Abbrev: PT, proximal tubule; TL, thin limb; TAL, thick ascending limb; DCT, distal convoluted tubule (DCT1 and DCT2); CNT, connecting tubule; CD, collecting duct (P, principal cells, IC, intercalated cells); Mes, mesangial cells; Endo, endothelial cells; SMC, smooth muscle cells; Fib, fibroblasts; Mac, macrophages; Mono, monocytes; DC, dendritic cells; DA-PT, damage-associated PT; DA-TAL, damage-associated TAL; DA-DCT, damage-associated DCT.

**Fig. S4.**
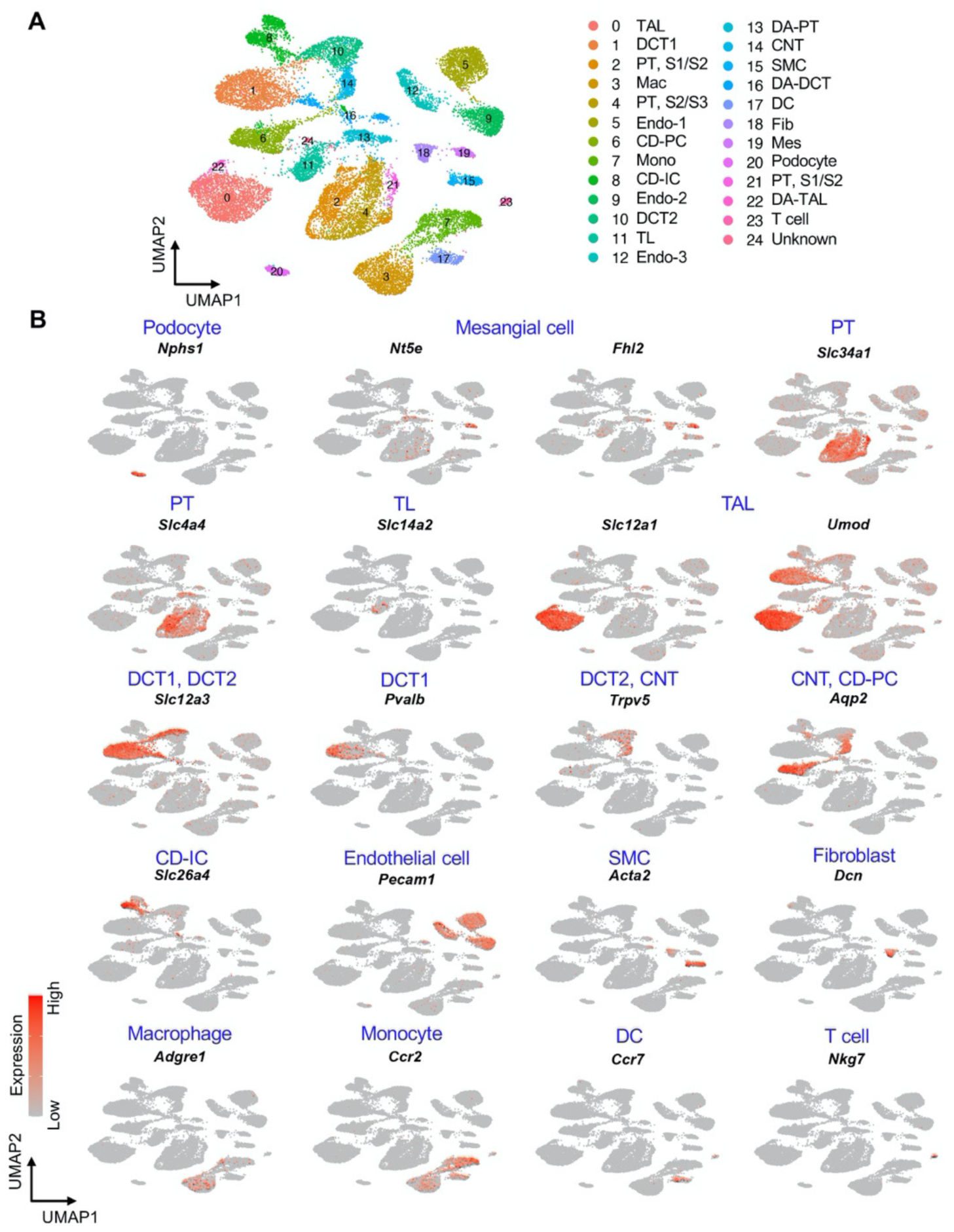
UMAP plots show the expression pattern of anchor genes in homeostatic and post-IRI kidneys. (**A**) UMAP plots show the identified cell clusters (resolution was set as 1.0). (**B**) UMAP plots show the expression pattern of indicated canonical marker genes (anchor genes) of each cluster. We manually combined 3 clusters of differentiated/mature proximal tubular cells (PT, S1/S2 and PT, S2/S3) into 1 cluster (PT) to generate a more coarse-grained cell-type annotation and data visualization in other figures. We also combined 3 clusters of endothelial cells (Endo-1, Endo-2, and Endo-3) into one cluster (Endo) for data visualization in other figures. S1, S1 segment; S2, S2 segment 2; S3, S3 segment of proximal tubule.

**Fig. S5.**
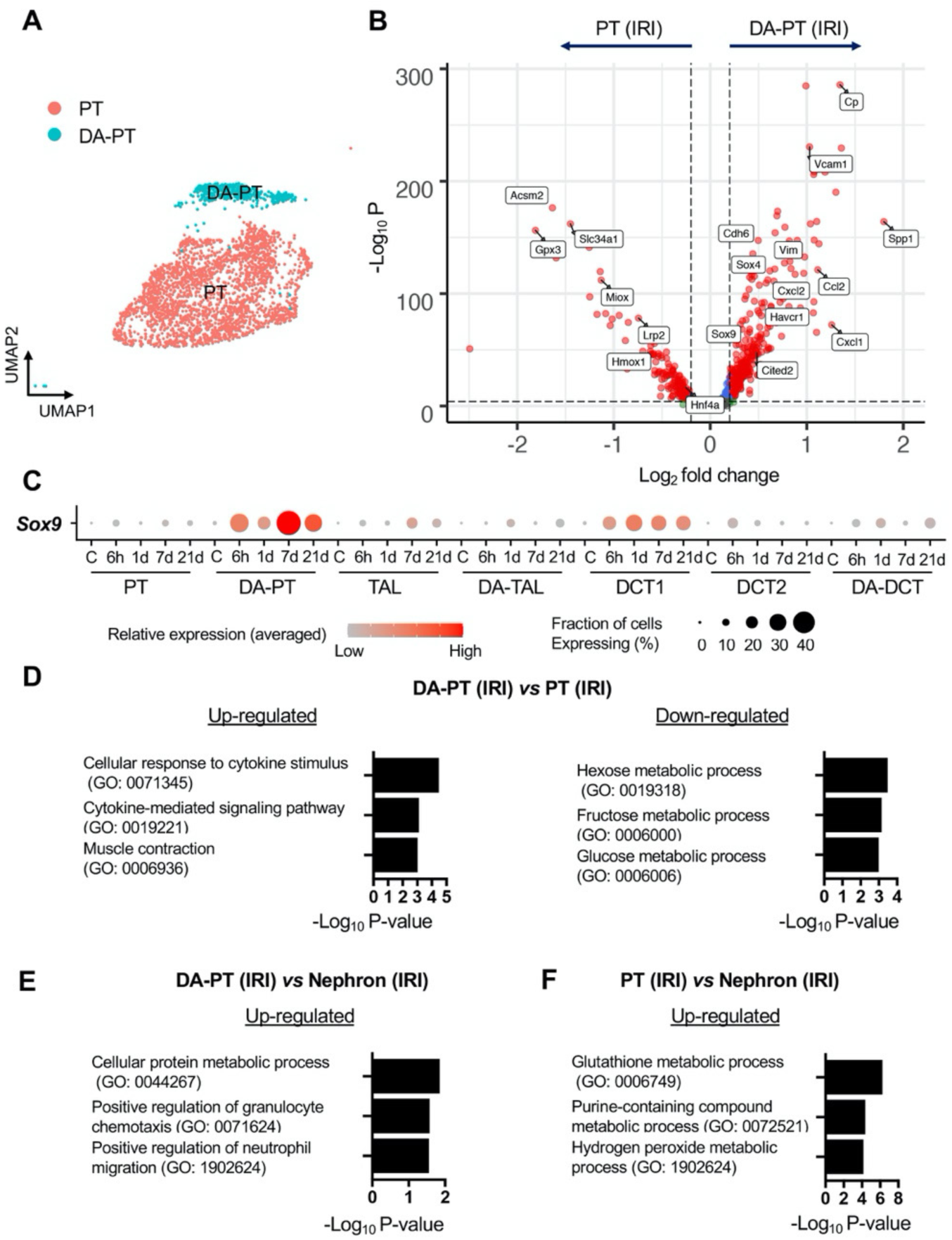
Damage-associated PT cells show an inflammatory transcriptional signature. (**A**) UMAP of PT clusters (PT, differentiated proximal tubular cell cluster; DA-PT, damage-associated proximal tubular cell cluster). See Fig. 1B. (**B**) Volcano plot showing a distinct transcriptional profile of damage-associated PT cells (PT cells in DA-PT cluster). A Wilcoxon rank-sum test was used for the statistical analysis comparing cells in PT cluster from IRI kidneys and cells in DA-PT cluster from IRI kidneys. Blue and grey data points indicate transcripts that fall below the set threshold for fold change (Log_2_ fold change, a threshold value of 0.25) and *P* value (a cutoff value of *10e-*5). Note that cells in the DA-PT cluster show reduced expression of homeostatic marker genes (*Acsm2*, *Slc34a1*, and *Lrp2*), oxidative stress defense genes (*Hmox*, *Miox*, *and Gpx3*), and a gene encodes transcription factor for PT maturation in renal development (*Hnf4a*). The cells in the DA-PT cluster also show enrichment of developmental genes (*Sox4*, *Sox9*, *Cited2*, and C*dh6*), damage-induced genes (*Havcr1* and *Vcam1*) and inflammatory cytokines and chemokines (*Spp1*, *Ccl2*, *Cxcl1*, and *Cxcl2*). (**C)** Dot plots show the expression of *Sox9* gene is highly enriched in damage-associated PT (DA-PT) cell population. **(D**) and (**E**) Gene ontology enrichment analyses identify that DA-PT cluster is enriched for immune response gene ontology. The cells in the DA-PT cluster from IRI kidneys (all time points) were compared with the cells in the PT cluster or cells in all other nephron segments and collecting duct (IRI kidneys, all time points) in (D) and (E), respectively. (**F**) Gene Ontology enrichment analyses identify the PT cluster is enriched for glutathione-mediated anti-oxidative stress responses. The cells in the PT cluster from IRI kidneys (all time points) were compared with the cells from all other nephron segments and collecting duct from IRI kidneys (all time points).

**Fig. S6.**
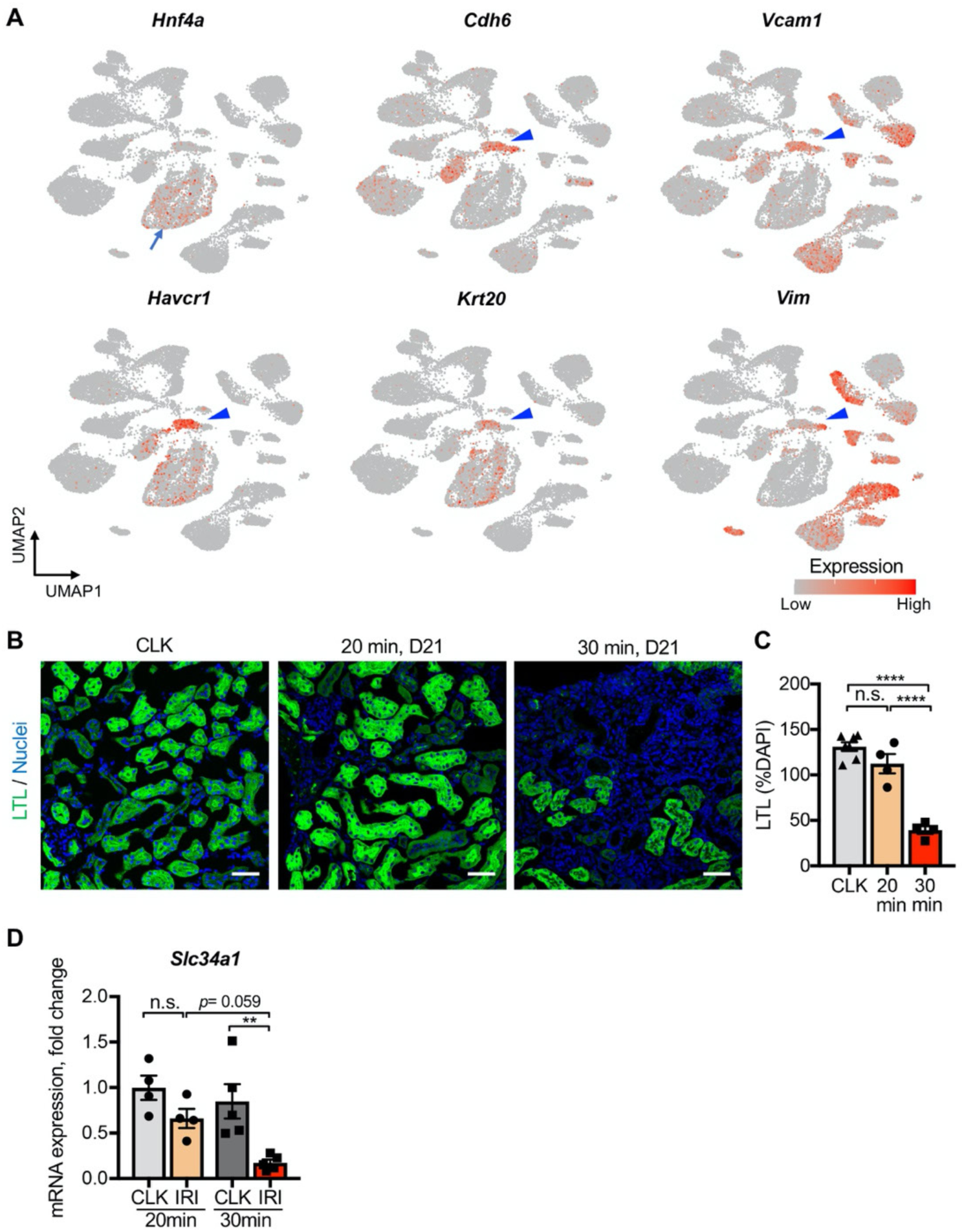
Severe IRI reduces expressions of proximal tubular differentiation markers. (**A**) UMAP plots showing the expression of indicated genes. Note that damage-associated PT cell state (cells in DA-PT cluster) are enriched with damage-induced genes (*Havcr1*, *Krt20*, *Vcam1*, and *Vim*) and exhibit less differentiated signature (upregulation of *Cdh6* and downregulation of *Hnf4a*). Arrow, PT cells; arrowheads; DA-PT cells. (**B**) and (**C**) Comparative analyses of kidneys underwent severe IRI (ischemic time 30 min) and mild IRI (ischemic time 20 min). Kidneys were harvested on day 21 post-IRI. Severe IRI reduces LTL^high^ proximal tubular cells. Lotus tetragonolobus lectin (LTL) binds fully differentiated proximal tubular cells. LTL^high^ areas from kidneys on day 21 were quantified. Wild-type C57BL/6J mice were used. Scale bars: 50 μm. N = 4-7. (**D**) Real-time PCR analyses of *Slc34a1*. mRNA expression of *Slc34a1* was reduced after severe IRI (ischemic time 30 min on day 21). Whole kidney lysates from post-IRI kidneys on day 21 were used. Contralateral kidneys (CLK) were used as controls. N = 4-5. \*\**P* < 0.01; \*\*\*\**P* < 0.0001, one-way ANOVA with post hoc multiple comparisons test.

**Fig. S7.**
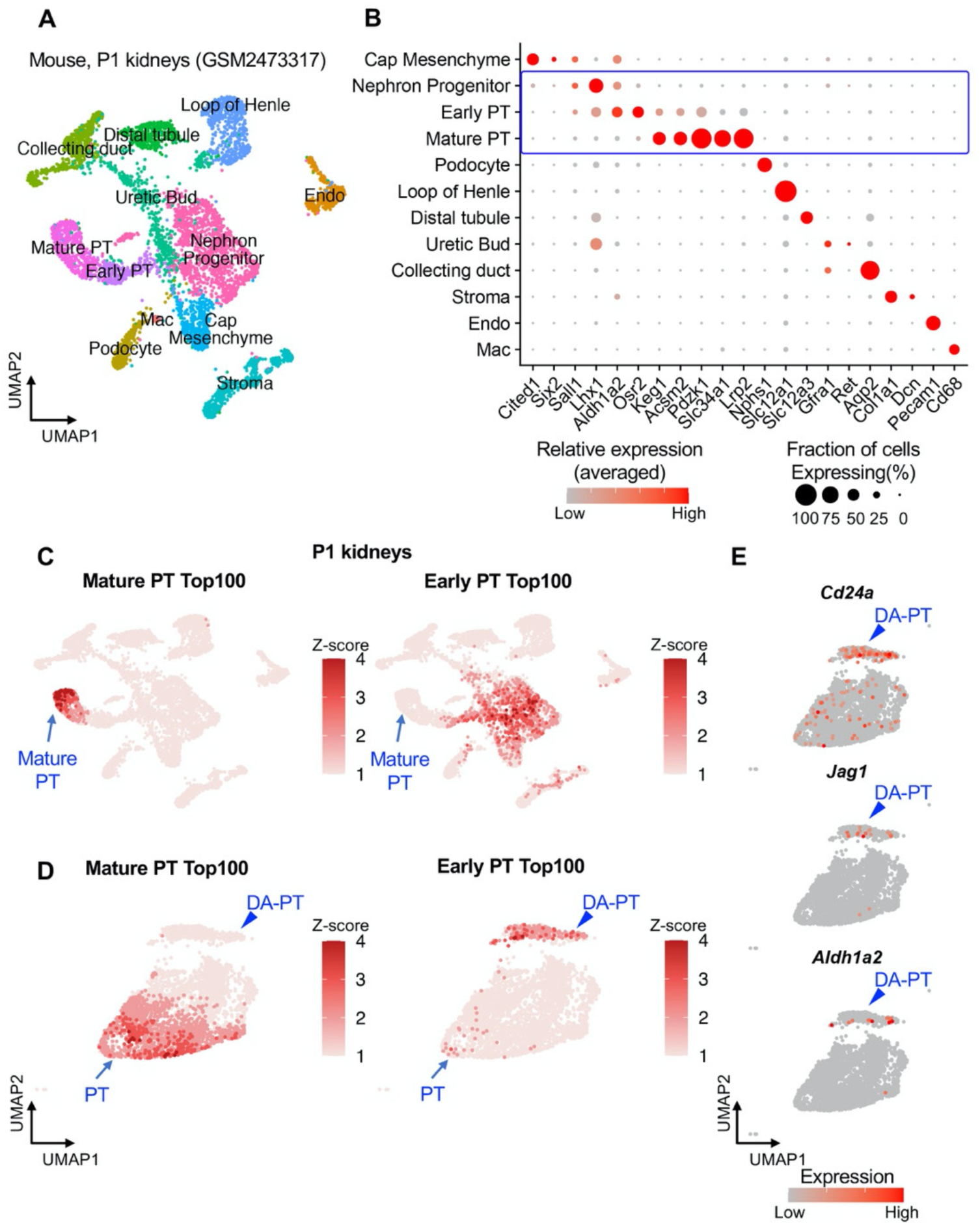
Comparative analyses of damage-associated PT cells and neonatal proximal tubular cells. **(A)** and (**B**) Characterization of mouse neonatal kidney single-cell RNA-seq data. UMAP plots show mouse neonatal kidney cells from GSM2473317 (4,693 cells; post-natal day 1) in (A). Dot plots show the gene expression patterns of cluster-enriched canonical markers in (B). (**C**) UMAP rendering of Top 100 genes characterizing mature and immature early proximal tubular (PT) cells. We obtained the top 100 genes representing mature PT and immature early PT cells by performing differential gene expression analysis using the “FindMarkers” command in Seurat. As expected, the mature PT Top 100 genes are enriched in mature PT cluster. The early PT Top 100 genes are enriched in immature early PT and nephron progenitor clusters. (**D**) UMAP rendering of mature and early PT genes on adult PT clusters from our dataset (PT and DA-PT). Note that the early PT Top 100 genes are highly enriched in damage-associated PT (DA-PT) population, indicating they are in a less differentiated state. (**E**) UMAP plots showing gene expressions of early PT genes (*Cd24a*, *Jag1*, and *Aldh1a2*) are enriched in DA-PT cluster.

**Fig. S8.**
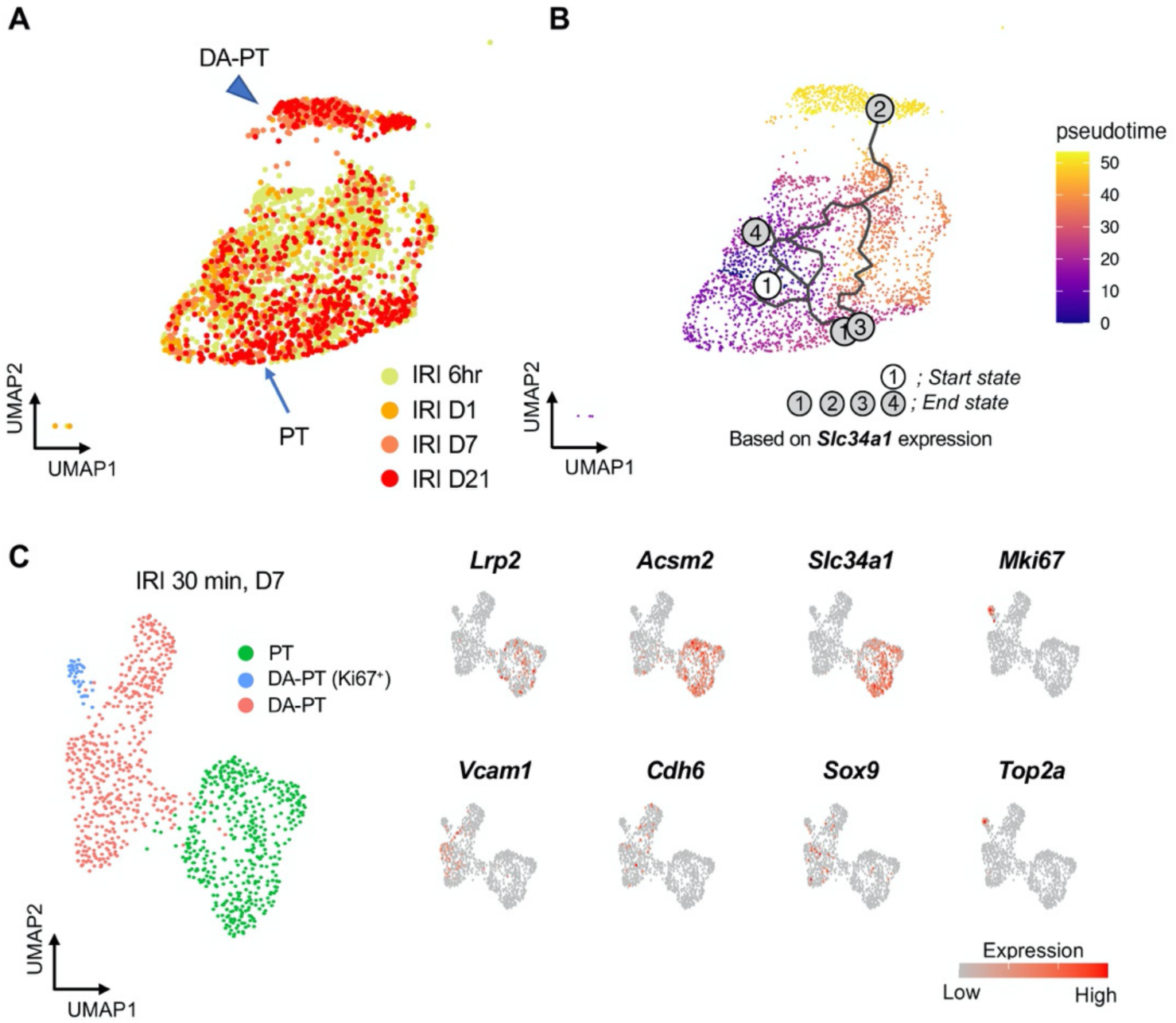
Trajectory analyses predict lineage hierarchy from differentiated mature PT cells to damage-associated PT cells. (**A**) UMAP plots showing mature and damage-associated PT cells (PT and DA-PT clusters) underwent IRI. (**B**) Pseudotime trajectory analysis using Monocle 3. A region occupied with *Slc34a1*^high^ cells was set as a starting state. Note that predicted trajectory starting from PT to DA-PT state. See Fig. 1F for the analysis with a different starting “root” setting. The earliest time point of injury was used as a starting root in Fig. 1F. Both analyses resulted in a similar predicted trajectory. (**C**) UMAP plots showing proximal tubular cells from PT and DA-PT clusters on IRI day 7 (D7). Cells from post-IRI kidneys on day 7 are shown and used for RNA velocity analysis to investigate potential cellular plasticity at this stage (single data point). See Fig. 1G for RNA velocities (trajectory). UMAP plots in the right panels show the indicated genes; homeostatic genes (*Lrp2*, *Acsm2*, and *Slc34a1*), damage-induced genes (*Vcam1*, *Cdh6*, and *Sox9*), and cellular proliferation (*Mki67* and *Top2a*). Fig. S8A and S8B are supporting data for Fig. 1F. Fig. S8C is supporting data for Fig. 1G.

**Fig. S9.**
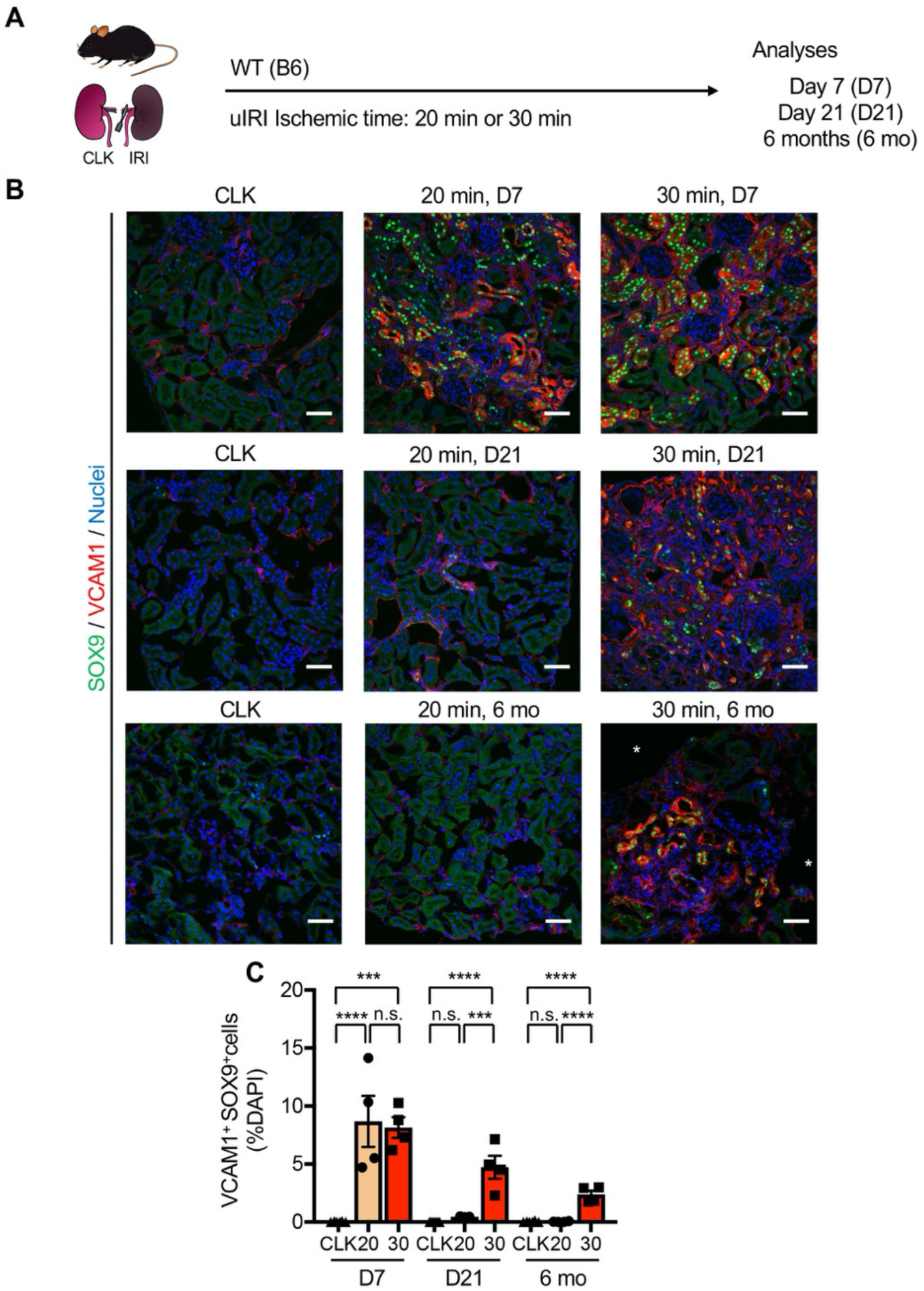
Comparative analyses of mild and severe IRI identify distinct temporal dynamics of damage-associated PT cells. (**A**) Experimental workflow. Wild type C57BL/6J mice were used. Mild IRI (20 min ischemia) and severe IRI (30 min ischemia). (**B**) Immunostaining for SOX9 and VCAM1. Note that double-positive cells (indicative for damage-associated PT cell state) emerge after mild and severe IRI similarly on day 7, but they are differentially regulated subsequently. SOX9^+^VCAM1^+^ cells disappear after mild IRI (20 min ischemia) on day 21, but they persist at least for 6 months after severe IRI (30 min ischemia). Scale bars: 50 μm. * indicates cystic lesions. (**C**) Quantification of double-positive cells over DAPI^+^ area in (B). N= 3-8. \*\*\**P* < 0.001, \*\*\*\**P* < 0.0001, one-way ANOVA with post hoc multiple comparisons test. n.s., not significant.

**Fig. S10.**
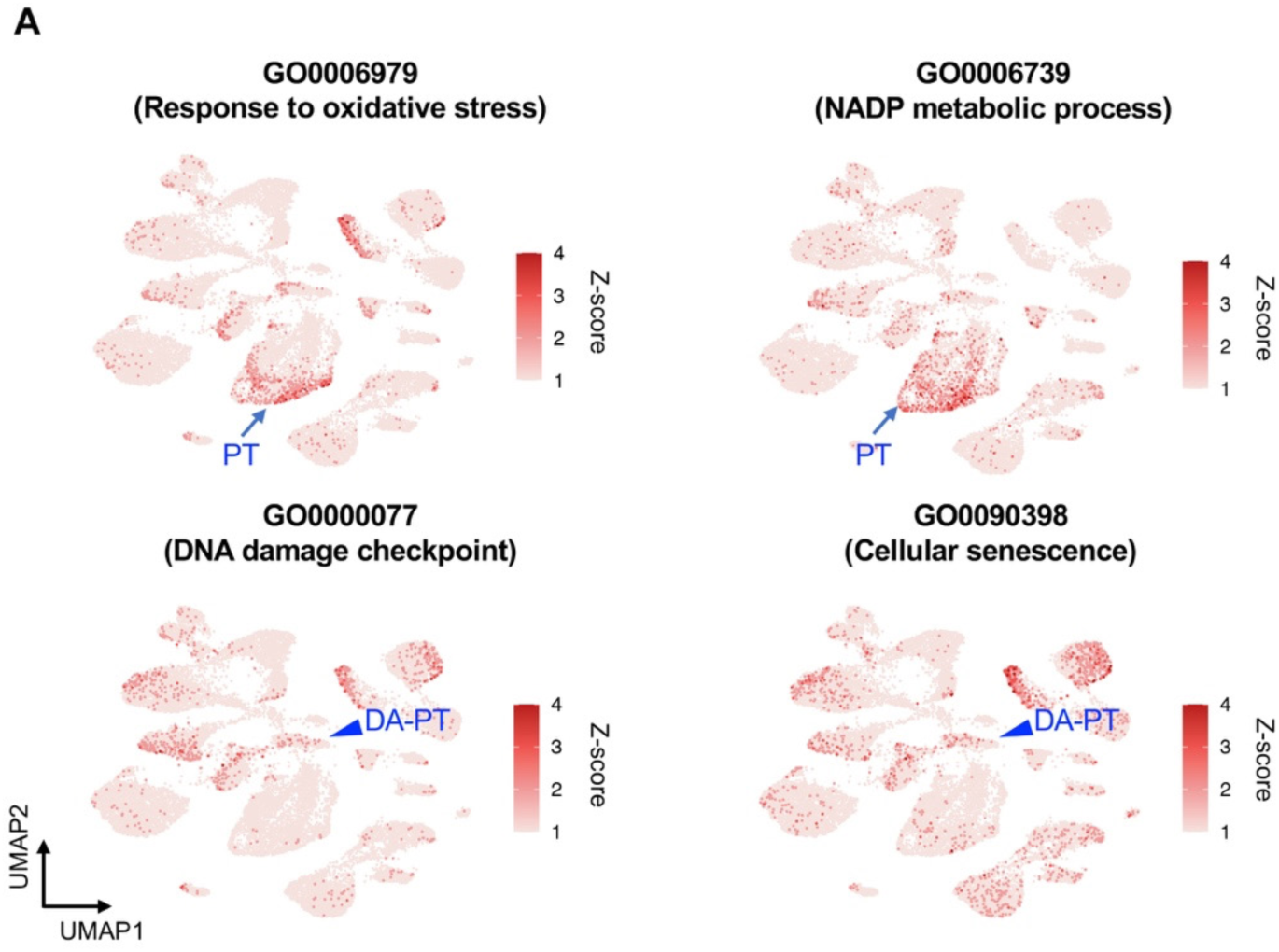
Gene ontology analyses identify enrichment of anti-oxidative stress defense genes in differentiated/mature PT cells. (**A**) UMAP rendering of signaling pathways. Upper panels show the pathways enriched in differentiated PT cells (PT cluster). Lower panels show the pathways enriched in damage-associated PT cells (DA-PT cluster). Arrows indicate differentiated PT cell cluster (PT). Arrowheads indicate DA-PT cell cluster.

**Fig. S11.**
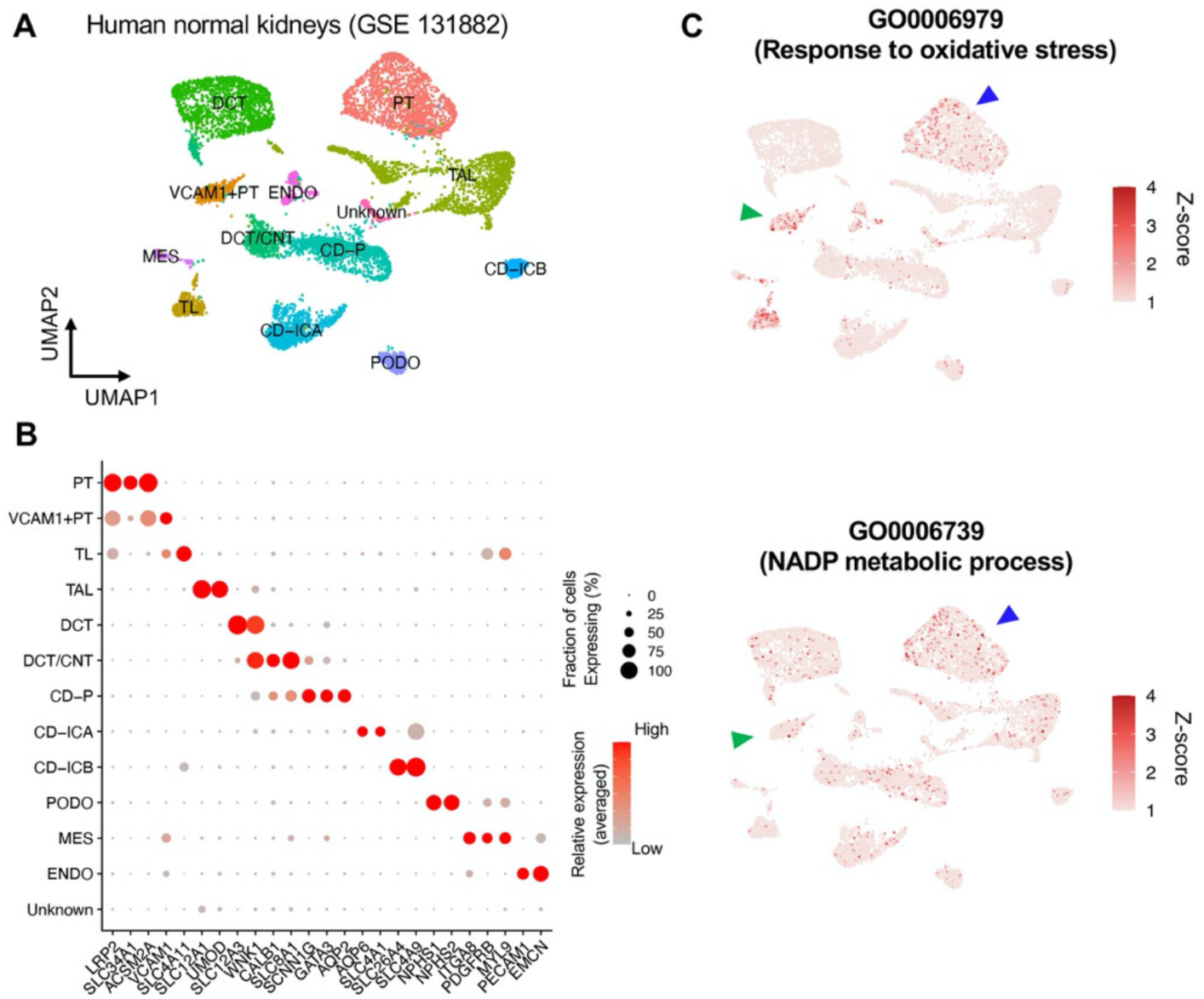
Characterization of human normal kidney single-nucleus RNA-seq data. (**A**) UMAP plots showing human normal kidney cells from GSE131882 (7,631 cells). Two normal kidney datasets were integrated and analyzed. (**B**) Dot plot showing the gene expression patterns of cluster-enriched canonical markers. (**C**) UMAP rendering of signaling pathways. Note that the signaling pathways for anti-oxidative stress, which are enriched in differentiated PT cells in mouse kidneys, are also enriched in normal differentiated PT cells in humans. Blue arrowheads: PT cells, green arrowheads, VCAM1^+^ PT cells.

**Fig. S12.**
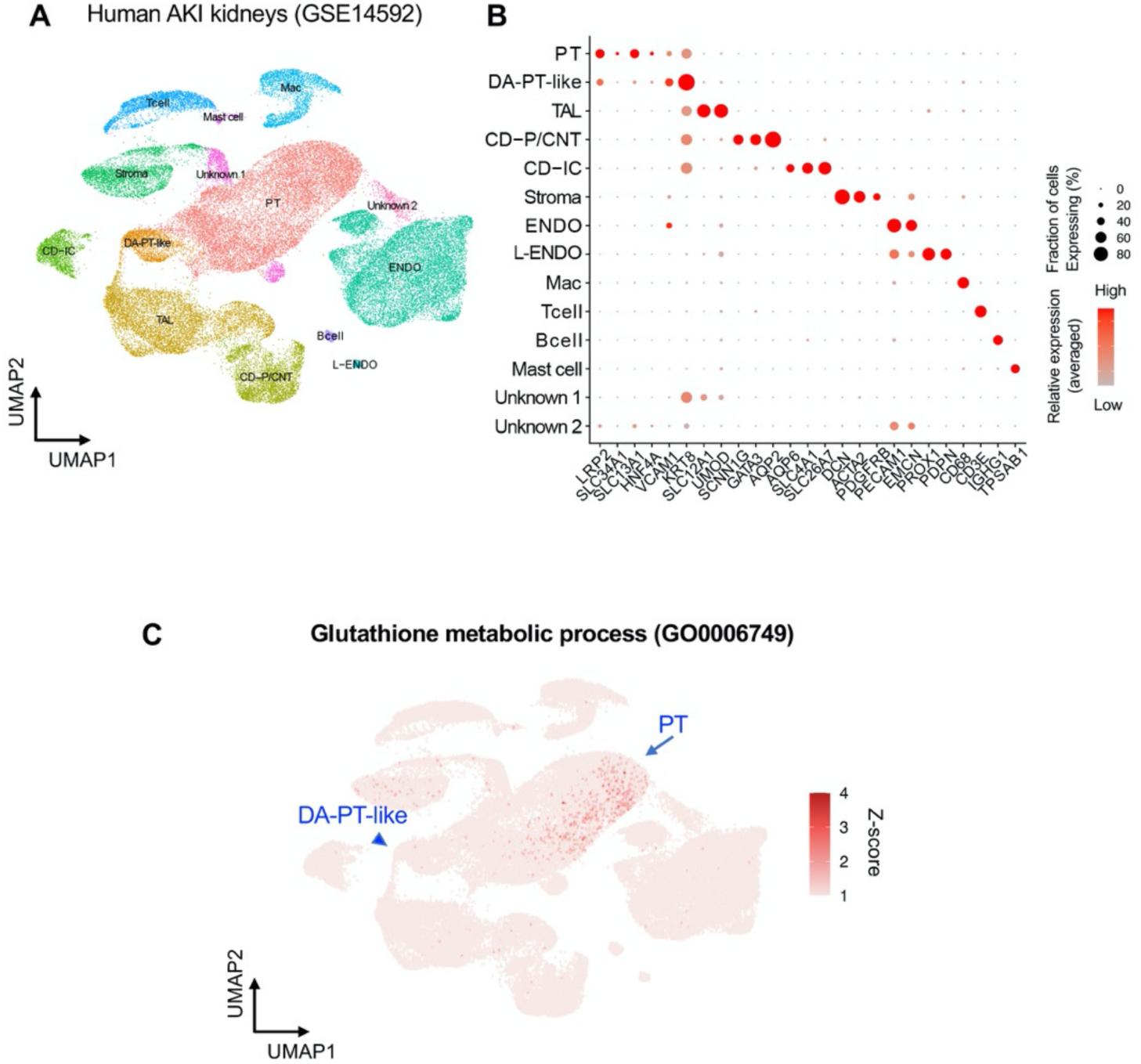
Characterization of human AKI kidney single-cell RNA-seq data. (**A**) UMAP plots showing human AKI kidney cells from GSE14592 (43,998 cells). Data from the two non-rejecting AKI tissues were analyzed and integrated. (**B**) Dot plot showing the gene expression patterns of cluster-enriched canonical markers. Note that DA-PT-like cells exhibit reduced expression of homeostatic genes (ex. *LRP2*, *SLC34A1*) but are enriched for damage-induced genes (ex. *VCAM1*, *KRT8*) as in the case of mouse damage-associated PT (DA-PT) cells. (**C**) UMAP rendering of signaling pathways. Note that genes for the glutathione-metabolic process are highly expressed in PT and gradually reduced towards DA-PT-like cells as in the predicted trajectory in Fig. 6F.

**Fig. S13.**
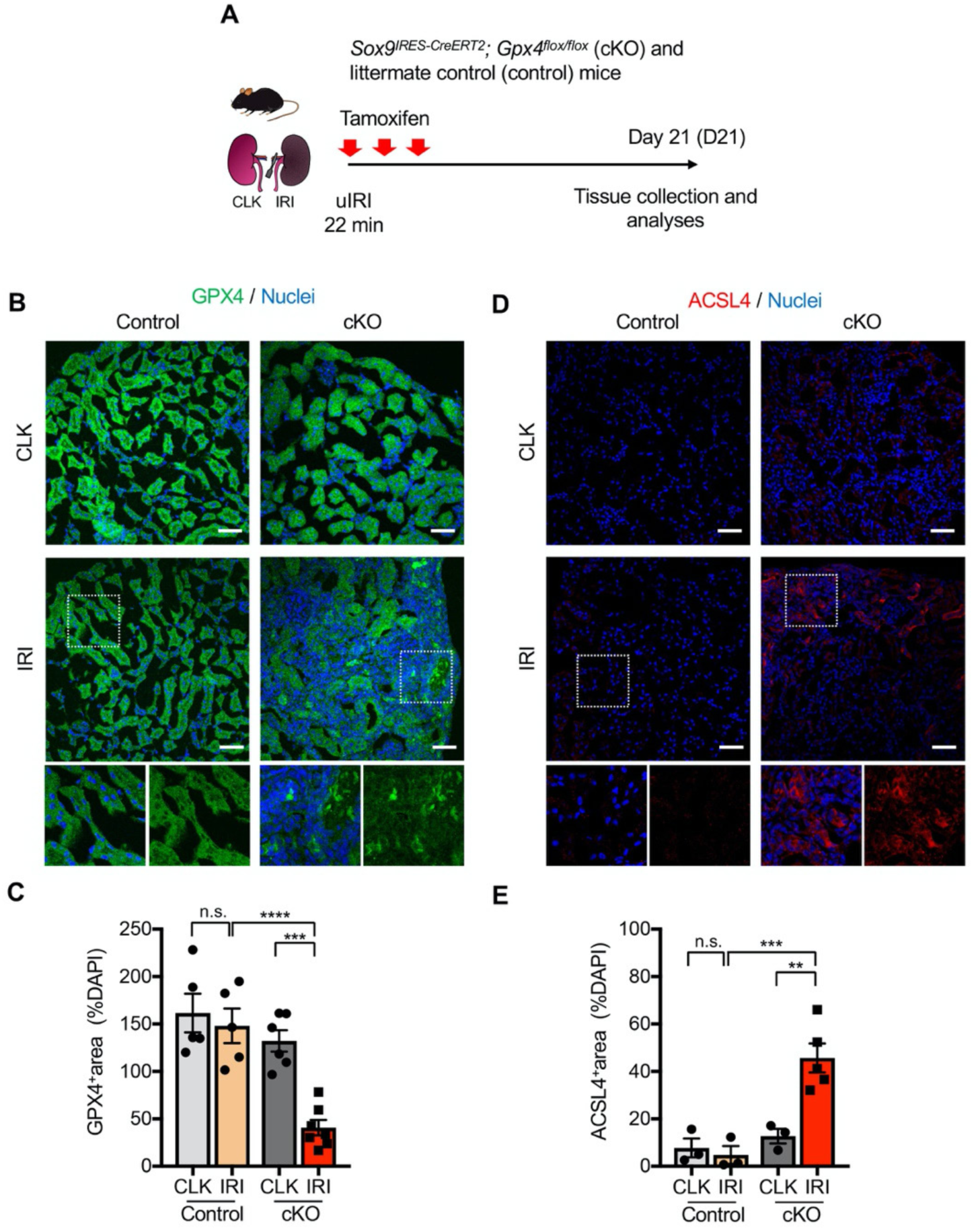
Genetic deletion of *Gpx4* leads to augmented ferroptotic stress after mild IRI. (**A**) Experimental workflow for genetic deletion of *Gpx4* in *Sox9*-lineage cells. cKO mice and their littermate controls were subjected to the same ischemic stress (ischemic time, 22 min) and tamoxifen treatment. Note that *Sox9* is only induced in proximal tubular cells after IRI (*Sox9* gene is silenced in this tubular segment after the completion of renal development); therefore, *Gpx4* deletion occurs after injury in IRI kidneys with tamoxifen injection. The initial dose of tamoxifen was administered immediately before the surgical intervention. Left kidneys were subjected to mild (22 min) ischemia. Only littermate controls were used for phenotypic comparisons. (**B**) and (**C**) Immunostaining for GPX4 and quantification. GPX4 immunostaining confirms the deletion of GPX4 in IRI-kidneys from cKO mice (See IRI, cKO). Inlets show the higher magnification and single-color images of dotted areas. N = 5-7. (**D**) and (**E**) Immunostaining for ACSL4 and quantification. Inlets show the higher magnification and single-color images of dotted areas. N = 3-5. \*\**P* < 0.01, \*\*\**P* < 0.001, \*\*\*\**P* < 0.0001, one-way ANOVA with post hoc multiple comparisons test. Scale bars: 50 μm in (B) and (D).

**Fig. S14.**
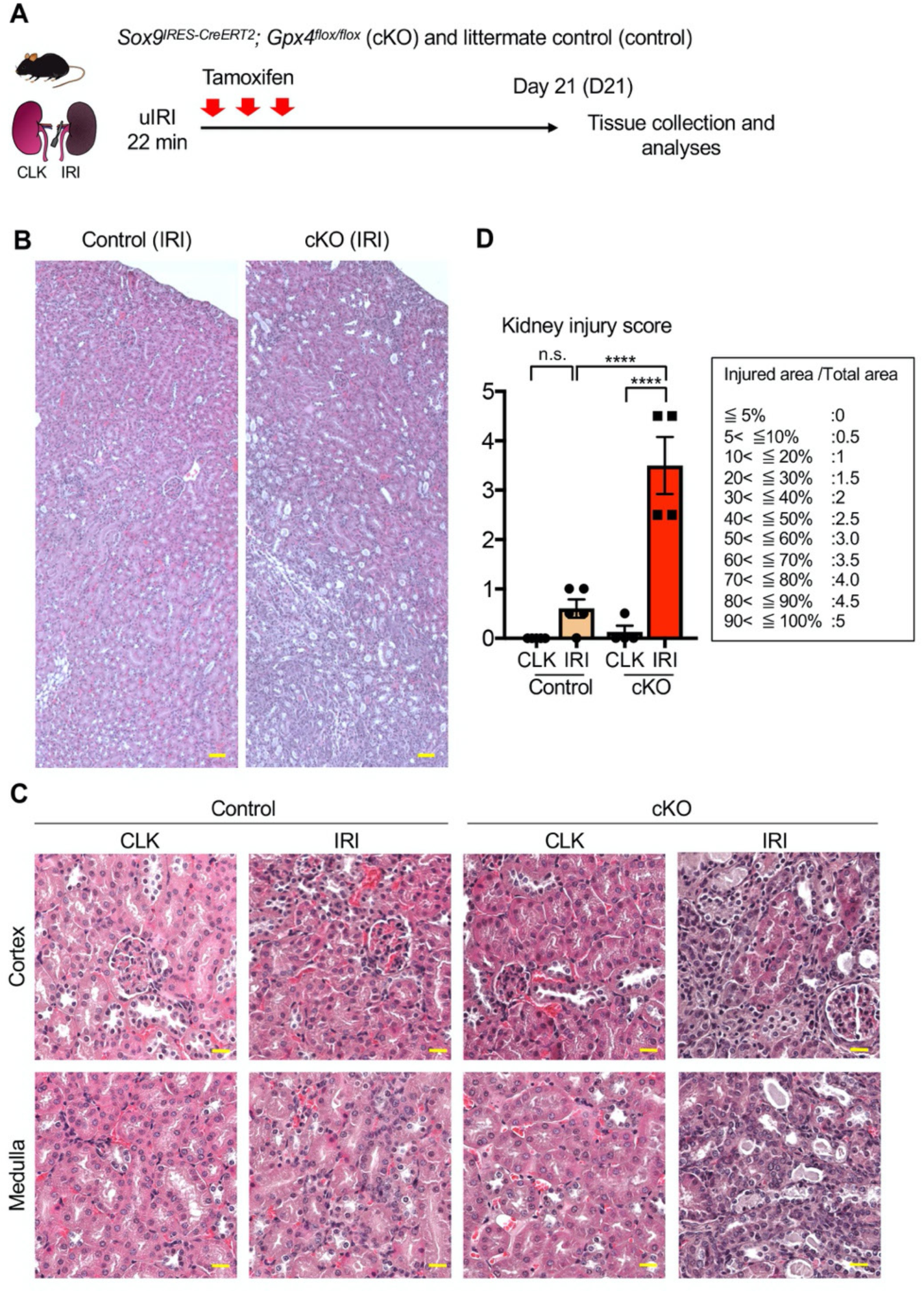
Genetic deletion of *Gpx4* leads to severe kidney injury after mild IRI. (**A**) Experimental workflow for genetic deletion of *Gpx4* in *Sox9*-lineage cells. cKO mice and their littermate controls were subjected to the same ischemic stress (ischemic time, 22 min) and tamoxifen treatment. (**B**) Hematoxylin-eosin staining of IRI kidneys. Control; littermate controls. Representative images from IRI kidneys are shown. (**C**) Higher magnification images. Note that *Gpx4* deletion after mild IRI resulted in severe tubular dilatation, flattening of tubular epithelial cells, cast formation, and inflammatory cell infiltration. (**D**) Quantification of kidney injury score. N = 4-5. \*\*\*\**P* < 0.0001, one-way ANOVA with post hoc multiple comparisons test. Scale bars: 50 μm in (B); 20 μm in (C).

**Fig. S15.**
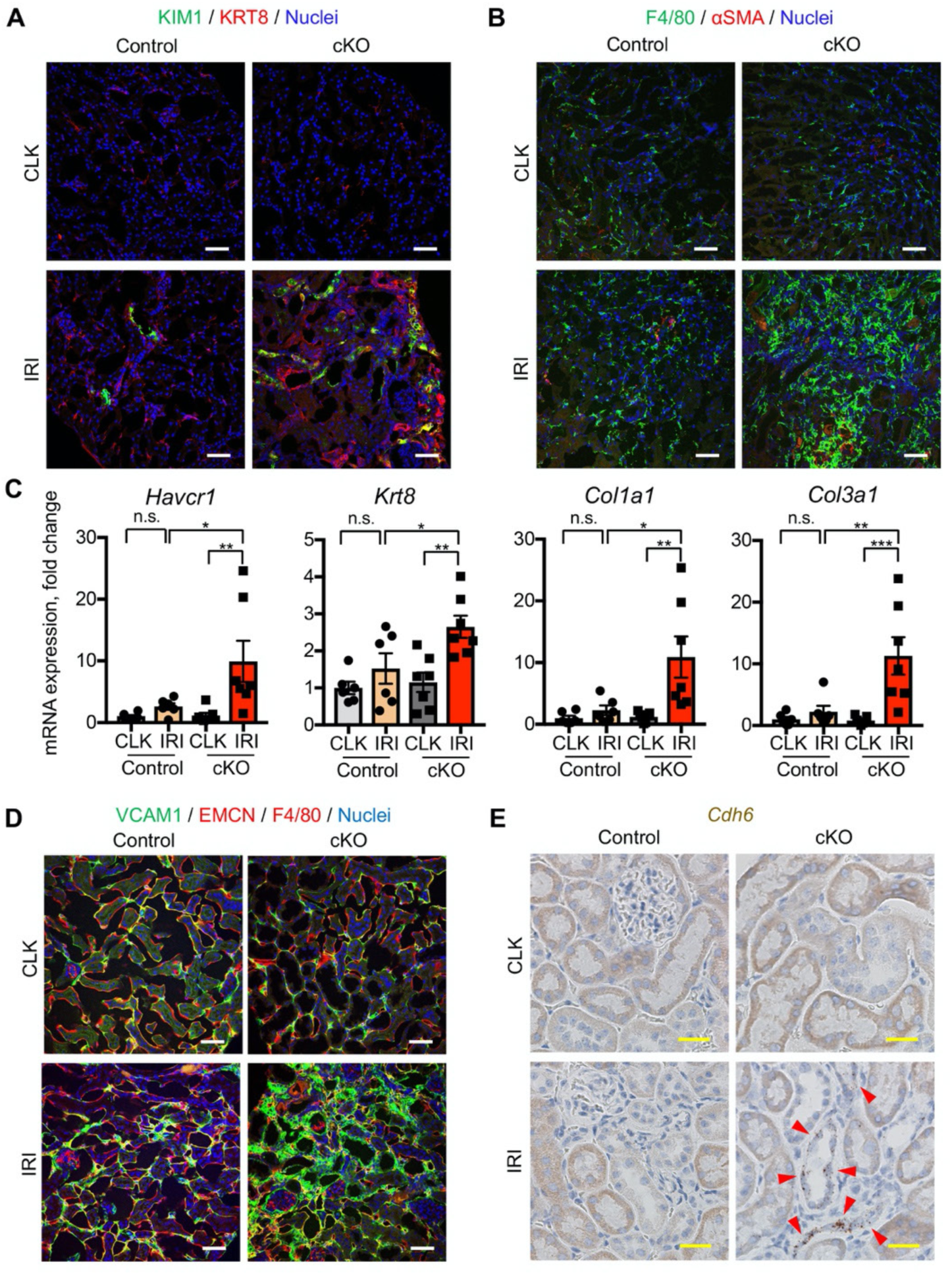
Genetic deletion of *Gpx4* leads to the accumulation of damage-associated PT cells. (**A**) and (**B**) Immunostaining for KIM1, KRT8, F4/80, and *α*SMA. Note that *Gpx4* deletion increased tubular injury markers (KIM1 and KRT8) and led to the accumulation of myeloid cells (F4/80) and myofibroblasts (*α*SMA) in post-IRI kidneys from cKO mice on day 21 (ischemic time, 22 min). (**C**) Real-time PCR analyses of indicated genes. N = 6–7. \**P* < 0.05; \*\**P* < 0.01, one way ANOVA with post-hoc multiple comparisons test. (**D**) Immunostaining for VCAM1, EMCN (endomucin), and F4/80. Note that VCAM1^+^EMCN^−^F4/80^−^ cells (damage-associated PT cell state) accumulated in *Gpx4*-deleted kidneys after mild IRI on day 21. (**E**) *In situ* hybridization for *Cdh6* expression. *Cdh6* is induced in tubular epithelial cells in *Gpx4*-deleted kidneys after mild IRI on day 21. Arrowheads: *Cdh6*+ tubular epithelial cells. These data indicate that damage-associated PT (DA-PT) cells accumulate even after mild IRI when the *Gpx4* gene is deleted. Note that littermate control kidneys underwent the same ischemic time (22 min) recovered without accumulating myofibroblasts, macrophages, and damage-associated PT cells, indicative of successful recovery. *Gpx4*-deleted kidneys underwent mild IRI (22 min ischemia) mimicked the failed renal repair phenotype observed after severe ischemic injury. Scale bars: 50 μm in (A), (B), and (D) and 20 μm in (E).

**Fig. S16.**
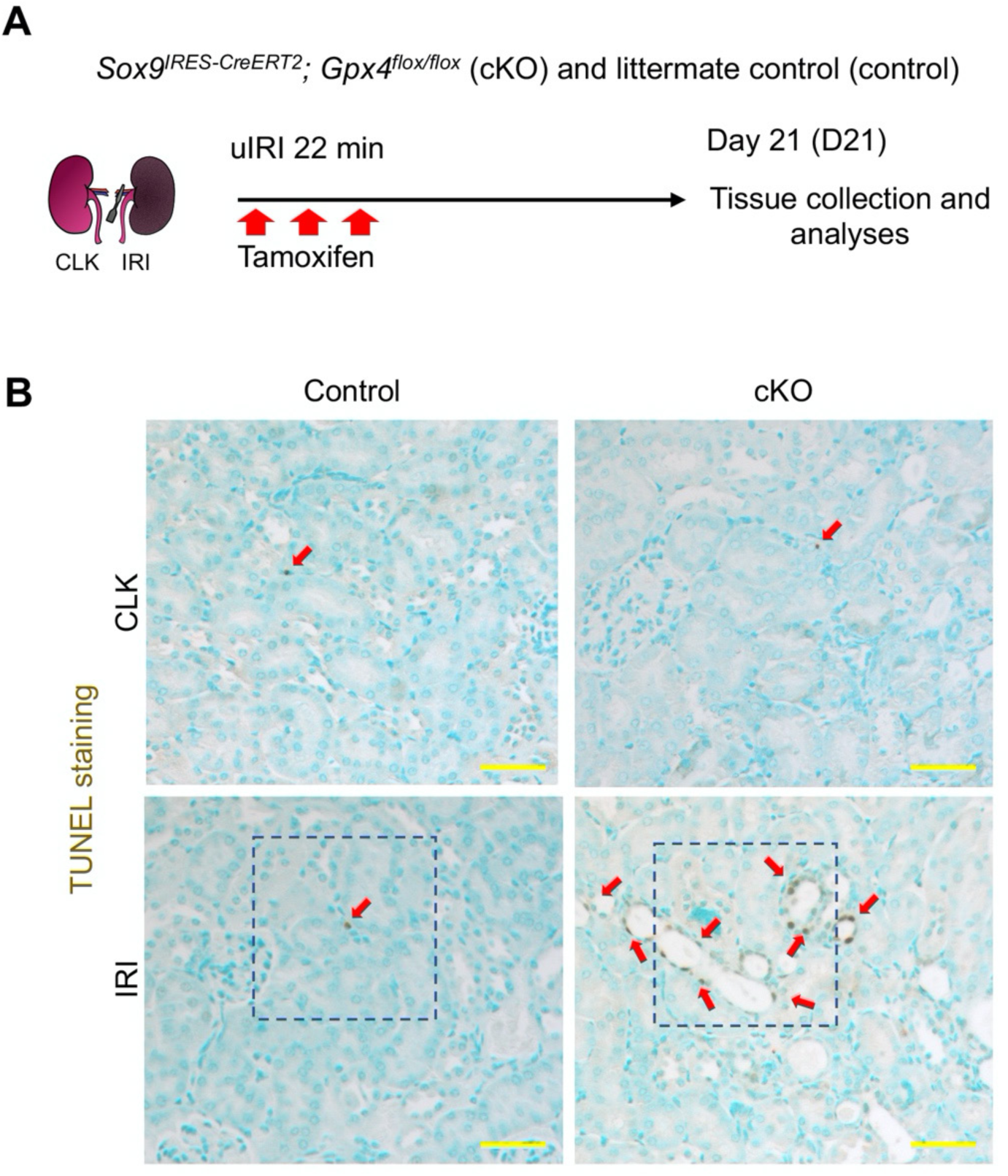
Genetic deletion of *Gpx4* leads to cell death of tubular epithelial cells. (**A**) Experimental workflow for genetic deletion of *Gpx4* in *Sox9*-lineage cells. cKO mice and their littermate controls were subjected to the same ischemic stress (ischemic time, 22 min) and tamoxifen treatment. (**B**) TUNEL staining for evaluating cell death. The dotted area are shown in Fig. 7G. See quantification for Fig. 7H. Scale bar, 50 μm. N=4. Arrows, TUNEL^+^ nuclei.

## Notes

**Competing interests:** The authors have declared no conflict of interest exists.

### Competing Interest Statement

The authors have declared no competing interest.

